# Inside-out Signalling From Aminopeptidase N (CD13) To Complement Receptor 3 (CR3, CD11b/CD18)

**DOI:** 10.1101/2021.12.28.474389

**Authors:** Laura Díaz-Alvarez, Mariana Esther Martínez-Sánchez, Eleanor Gray, Enrique Ortega

**Author notes:** This article is dedicated to the memory of Dr. Heliodoro Celis Sandoval, friend and mentor who will be dearly missed.

## Abstract

Upon ligand engagement, certain receptors can activate an integrin through a mechanism called inside-out signalling. This phenomenon prepares the cell for the next steps of the process it will perform. CR3 (Complement receptor 3), the most abundant β2 integrin in monocytes and macrophages, and CD13 (aminopeptidase N) are two immune receptors with overlapping activities: adhesion, phagocytosis of opsonized particles, and respiratory burst induction. They can be found together in functional signalling microdomains, or lipid rafts, on the surface of human leukocytes. Thus, given their common functions, shared physical location and the fact that some phagocytic and adhesion receptors activate a selection of integrins, we hypothesized that CD13 could activate CR3 through an inside-out signalling mechanism. To test this hypothesis, we first ascertained the activation of CR3 after CD13 crosslinking in human monocyte-derived macrophages. We used an integrated analysis of bioinformatics and experimental data to suggest two possible signalling cascades that could explain the phenomenon. Finally, we show that the non-receptor tyrosine kinase Syk is a key attenuator of this pathway. Our results demonstrated that, even in the absence of canonical signalling motifs, and despite having a noticeably short cytoplasmic tail (7-10 amino acids), CD13 was capable of triggering an inside-out signalling cascade, adding a new function to those already known for this moonlighting protein.

**One Sentence Summary:** Stimulation of CD13 activated the integrin CR3 via an inside-out signalling pathway, a mechanistic model is proposed.

## INTRODUCTION

CD13 is a cell-membrane ectoenzyme (E.C.3.4.11.2) also known as aminopeptidase N due to its capacity to cleave N-terminal (preferentially neutral) amino acid residues (aa) from peptides, CD13 enzymatic activity is Zn^2+^-dependent; thus it is considered to be a metalloproteinase. CD13 is expressed in a number of cell types such as several epithelia, myeloid cells and during the early stages of lymphocyte differentiation (*1*). Most of its 960 aa are located extracellularly; roughly 25 aa constitute the transmembrane portion and only 7-10 aa correspond to the intracellular portion of the protein (*2*). The intracellular and extracellular segments of CD13 have distinct functions. The enzymatic activity is located on the extracellular domains, and accounts for CD13’s role in the processing of bioactive peptides. The intracellular portion, on the other hand, is able to mediate signal transduction when the receptor is crosslinked in a peptidase activity-independent fashion. In this way, CD13 mediates cellular processes like phagocytosis, cell migration and adhesion (*3–5*). Signal transduction takes place despite the shortness of the intracellular tail with only, the presence of a single potential p-Tyr and the absence of classical signalling sequences like ITAMs. Due to the range activities of CD13, it is considered as a “moonlighting” protein.

Complement receptor 3 (CR3) is a member of a group of membrane proteins called α/β integrins. These are cell surface molecules that have functions related to cell-cell and cell-extracellular matrix (ECM) adhesion. Additionally, some integrins mediate respiratory burst, and phagocytosis. The α/β integrins are heterodimers comprised of one α and one β chain. There are 18 mammalian genes coding α chains and 8 coding β chains that form the 24 α/β integrins reported in vertebrates (*6*). Integrins are grouped based on their β chains. Heterodimers with the constant β chain is CD18 are called β2 integrins; a family of four members that is expressed in leukocytes, where the variable a chain is either αL, αM, αX or αD, also referred to as CD11a, b, c, or d, respectively. CR3 corresponds to the CD11b/CD18 combination and is also known as Mac-1 and integrin αM/β2. The CD11b and CD18 subunits are 155-165 kDa and 94 kDa polypeptides, respectively (*7, 8*).

CR3 is primarily expressed in leukocytes like neutrophils, monocytes, macrophages, and dendritic cells (*9*). Two main physiological roles have been described for CR3. Firstly, it acts as a phagocytic receptor for particles and pathogens opsonized with iC3b complement fragments. Such pathogens are engulfed and subsequently destroyed by the phagocytes (reviewed in (*10*)). Secondly, CR3 is an adhesion molecule that participates in leukocyte extravasation during inflammation. This is due to its ability to bind ligands present on endothelial cells, such as ICAM-1, ICAM-2, JAM-A, JAM-C, and RAGE (*11, 12*).

Integrins can exist in three conformations: open, closed, and intermediate. These are causally related to ligand affinity, that is, high, low, and intermediate affinities, correspondingly. The larger and best-known of their integrin ligands like fibrinogen, collagen, and fibronectin, can only bind integrins in the open conformation. Thus, this conformation can be considered to be the fully activated state of the receptor. However, smaller ligands and peptides bearing the sequence Arg-Gly-Asp can interact with the closed receptor. The intermediate-affinity state can be regarded as an additional inactive state since the extracellular ligand-binding site is not exposed, however, intracellular signals can interact with the receptor in this conformation (reviewed in (*13, 14*)).

Immune receptors like CD13 and CR3 do not operate as isolated entities, they are part of an array of molecules and intracellular signalling events that act in concert to promote cellular functions. Upon ligand engagement by a receptor, two different types of signal transduction events can be distinguished. The first is called outside-in signalling, where an external signal is transmitted to the inside of the cell. This process is mediated by the sequential activation of intracellular molecules such as tyrosine kinases, GTPases, nuclear factors, etc, that culminate in a cellular function like gene expression, release of soluble mediators, polymerization of the actin cytoskeleton, and others. The second type of event is known as inside-out signalling, where an internal signal changes the conformation of receptors that face the outside of the cell. This process also involves the stimulation of a series of tyrosine kinases and phosphatases, GDP-GTP exchanging factors, etc, but in this case the pathway results in the activation of a different cellular receptor. Inside-out signalling can be seen as the pre-activation of the surface molecules required for the next stages of a cellular process. For example, polymorphonuclear leukocytes in circulation must slow down prior to its firm adhesion to the endothelium and extravasation. During the initial interaction of leukocytes with endothelial cells, chemokines and selectins binding to their receptors on the endothelium activate integrins like LFA-1, CR3 and VLA-4 via inside-out signalling, which launches the next phase of extravasation: firm adhesion (*15*).

The activation of CR3, that is the transition from its low affinity to its high affinity conformations, occurs either through ligand recognition (outside-in signalling) or via an intracellular signal coming from a different cell surface receptor (inside-out signalling). The events triggered by either signalling pathway generally recruit distinct effectors, for example, following engagement of the extracellular effector Mindin, CR3 outside-in signalling results in the activation of the MAPK pathway and translocation of NF-κb into the nucleus (*16*). In contrast, stimulation of G-protein coupled receptors results in the transition of CR3 from its low-affinity to its high-affinity state via an inside-out signalling pathway that includes the activation of phospholipases Cβ2 and Cβ3, and the calcium- and diacylglycerol-regulated guanine nucleotide exchange factor I (reviewed in (*9*)). Some molecules can participate in both CR3 inside-out and outside-in signalling, including Rap1, RIAM, Talin, Kindlin and Syk (*9, 17–20*).

Syk (Spleen tyrosine kinase) is a 72 kDa non-receptor tyrosine kinase containing two SH2 domains and an active site-bearing domain. It is a key player in the immune system, orchestrating a wide range of responses such as FcγR-mediated phagocytosis, and BCR activation, (reviewed in (*21, 22*). The canonical mechanism for Syk activation involves a post-stimulation conformational change that allows Tyr-containing sequences called ITAMs on the receptor, to be phosphorylated by a protein kinase of the Src family. This creates a docking site to which Syk binds through its SH2 domains, cousing Syk to be activated by auto- phosphorylation. Once detached from the ITAM, Syk continues to phosphorylate its downstream substrates, until phosphatases like SHP-1 and SHP-2, deactivate it and signal transduction stops, (reviewed in (*23*)). Syk participates in a variety of signalling cascades, including those induced by the moonlighting ectoenzymes CD38 and CD13 (*3, 24*).

Our group previously found that upon CD13 stimulation, Syk is phosphorylated and the adaptor molecule Grab2 and the Ras-GEF Sos1 co-precipitate with the receptor (*3, 25*). Additionally, research from our laboratory and others demonstrated that CD13 crosslinking by monoclonal antibodies (mAbs) enhances cellular functions mediated by other receptors such as FcγRs, other matrix metalloproteases and, most likely, scavenger receptors (*26–28*). For example, Mina-Osorio and Ortega (*26*) showed that co-crosslinking of CD13 and CD64 (FcγRI) synergistically augments both the efficiency of phagocytosis, and the duration of Syk phosphorylation in human monocytic cells.

In summary: i) integrins like CR3 can be activated by engagement of other receptors through a mechanism known as inside-out signalling, and ii) CD13 and CR3 share some of their mediated functions (phagocytosis, adhesion, and respiratory burst). Moreover, CD13 and CR3 can be found in physical proximity as both are present in lipid rafts (*29*), which is a strong indicator of a functional relationship. Finally, a link between the surface expression of CD13 and a different integrin (αvβ3) has already been recognized in breast cancer (*30*). In this work we report the possibility of the existence of a signalling pathway that links CR3 and CD13 in human monocyte-derived macrophages (MDMs), employing an integrated analysis of bioinformatics and experimental data.

First, we ascertained that CD13 crosslinking by antibodies causes the activation of CR3. Second, we establish that Syk is an attenuator of the inside-out signalling cascade initiated by CD13 crosslinking. Third, we interrogated molecular ontology bioinformatic databases, ran text mining analyses and manually curated a functional protein interaction network to suggest the components of the CD13-CR3 signal transduction pathway. Our findings have implications for the study of conditions in which the expression of CD13 is related to disease progression, as it is in breast cancer, where CD13 is related to the development of metastases (*31*), a phenomenon largely driven by integrins.

## RESULTS

### CD13 crosslinking resulted in the activation of CR3 (CD11b/CD18)

CR3 can exist in two main conformational states that correspond to a high or low affinity for its ligands, the active and inactive states, respectively. The high affinity state can be reached either by encountering its ligand (outside-in signalling) or by cell stimulation through other immune receptors, i.e. by inside-out signalling. Since CD13 crosslinking by antibodies increases the function of several receptors including those allowing the phagocytosis of zymosan, *E. coli* and IgG-opsonised particles (*3, 27*), we hypothesized that it could also promote the activation of integrins like CR3.

For this reason, we assessed the activation status of CR3 (CD11b/CD18) following CD13 crosslinking on human MDMs. CD13 molecules on the surface of MDMs were crosslinked using Fab fragments of the anti-CD13 antibody mAb C (Fab C) as primary antibody and, GαM F(ab)’2 fragments as secondary antibody. Next, cells were stained with a FITC-anti-CD11b(activated) antibody and analysed in the flow cytometer. Cells were first gated for size and granularity (Fig. 1A), then for singlets (Fig. 1B) and finally for median fluorescence intensity in the BL1 (FITC) channel (Fig. 1C and 1D). Fig. 1C shows the controls, i.e., unstained cells as well as cells incubated either without primary and secondary antibodies (No Fab) or with only secondary antibody (No Fab + secondary) followed by staining with anti-CD11b(activated) antibody. The resulting histograms demonstrate that incubation in the absence of Fab C does not produce a nonspecific anti-CD11b(activated) signal. In contrast, in panel D it is possible to distinguish CR3 activation in a representative sample of MDMs incubated with primary (Fab C) and secondary antibodies, i.e. after CD13-crosslinking. Fig. 1E shows the average and standard deviation of CD11b activation in CD13-crosslinked cells from a series of independent experiments. An average 43% (±16.4) of cells showed CD11b activation after CD13 crosslinking, while controls showed no activation. A one-way ANOVA followed by a Dunnett’s multiple comparisons test confirms that the difference between samples and controls was statistically significant at 95% (Dunnett’s CI95% -54 to -31). We ruled out the possibility that CR3 activation was not detected in all CD13 crosslinked cells due an incomplete occupation by Fab C. Fig. 1F shows a representative histogram from MDMs incubated with Fab C and a secondary antibody coupled to FITC, showing that Fab C bound efficiently to all cells.

**Fig 1.**
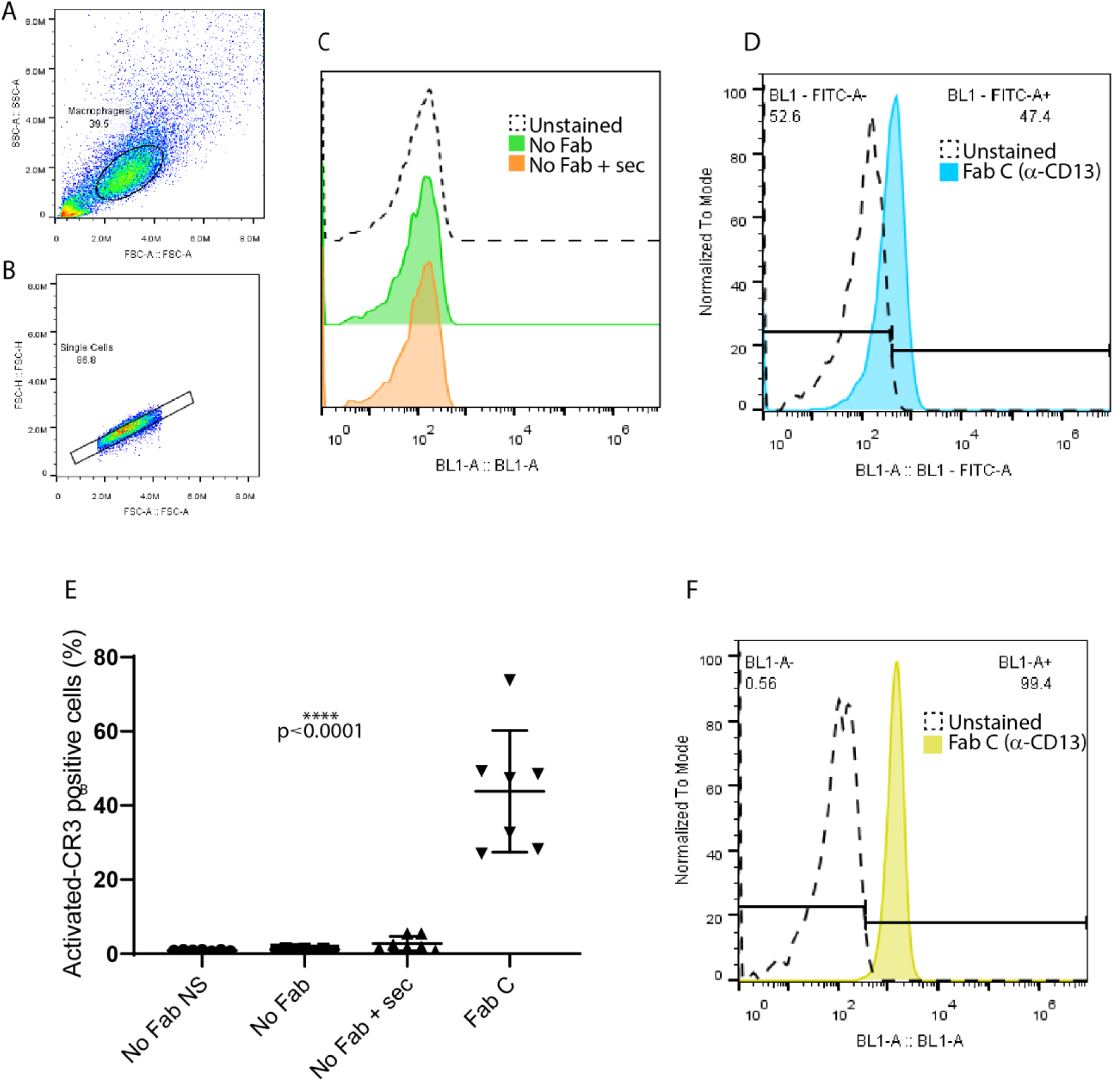
CD13 crosslinking activates CR3 in MDMs. Cells were first gated for size and granularity (**A**), then for singlets (B) and finally for median fluorescence intensity in the BL1 (FITC) channel (C and D). Controls are shown in panel C. In panel D a representative histogram of CR3 activation is shown from a 3h-incubation sample crosslinked with Fab C (anti-CD13) and secondary antibody F(ab)’2 fragments vs the unstained control. Panel E shows the average and standard deviations from 7 independent experiments. An average of 43.8% (±16.4) of cells showed CR3 activation after CD13 crosslinking. A one-way ANOVA followed by a Dunnett’s multiple comparisons test confirms that the difference between samples and controls is statistically significant, for both tests p<0.0001 (CI 95% -54 to -31.8). In panel F a representative histogram is shown, demonstrating that virtually all cells are positive for the CD13 stain.

CD13 crosslinking-dependent CR3 activation was specific for CD13, as crosslinking a different receptor (CD32) with anti-CD32 antibody fragments (Fab IV.3) did not elicit the same effect. Here CD32-crosslinked cells were treated and analysed in the same way as CD13-crosslinked cells. Fig. 2A and 2B show the gating process, first by size and granularity, then by the detection of singlets. The selected events were examined in the BL1 (FITC) channel as the anti-CD11b (activated) antibody was coupled to FITC. Fig. 2C displays representative histograms from CD13- and CD32-crosslinked cells, and their controls, where, as expected, only CD13- crosslinked MDMs were positive for the binding of the CD11b (activated) antibody. To validate this measurement, we repeated the assay twice more (Fig. 2D) and performed a one-way ANOVA followed by a Dunnett’s multiple comparisons test that confirmed the statistical significance of these results (Dunnett’s CI 95% -32 to -16). The lack of CR3 activation in cells incubated with Fab IV.3 was nor due to poor binding of this Fab to the cells, as Fig. 2E shows that all cells were positive for Fab VI.3 binding. Therefore, our results confirmed that crosslinking CD13 on human MDMs induced the activation of CR3 in a specific fashion.

**Fig. 2.**
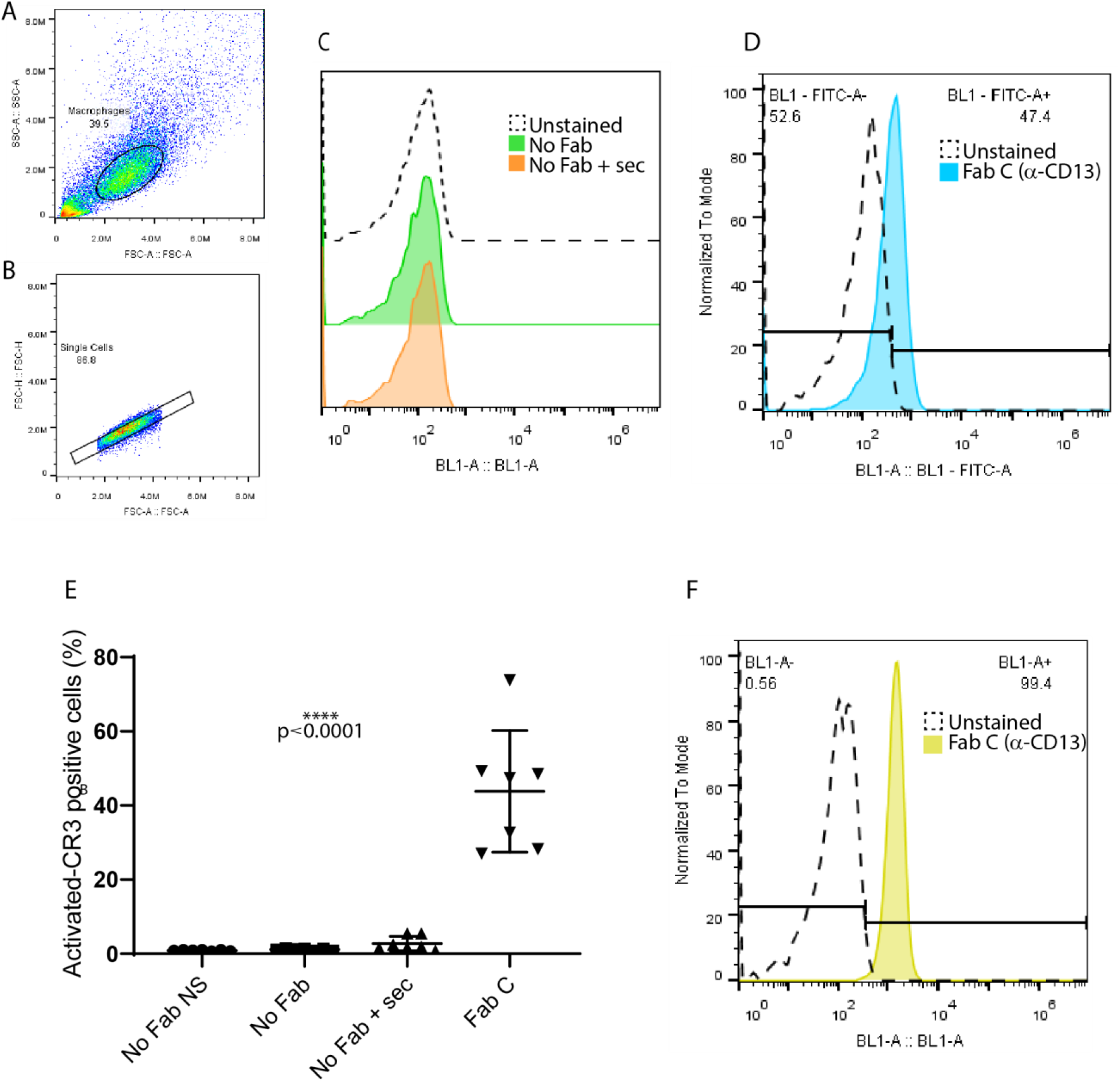
Anti-CD32 crosslinking does not induce CR3 activation in MDMs. In order to rule out the possibility that the activation of CR3 occurs as a consequence of the crosslinking of any immune receptor, MDMs were incubated in serum-free RPMI and CD13 or CD32 on their surface were crosslinked. Cells incubated without Fab were used as unstained control. Cells were first gated for size and granularity (**A**), then for singlets (B) and finally for median fluorescence intensity in the BL1 (FITC) channel (C). In the histograms from a representative experiment (panel C) it is possible to see that, unlike Fab C (anti-CD13), Fab IV.3 (anti-CD32) does not induce CR3 activation. Panel D shows the average percentage of CR3-activated cells and standard deviations from 3 independent experiments. A one-way ANOVA followed by a Dunnett’s multiple comparisons test confirms that the difference between CD13-crosslinked samples and the rest of the conditions is statistically significant, for both tests p<0.0001 (CI 95% -32.7 to -16.3). Inefficient Fab IV.3 binding is not responsible for the lack of CR3 activation since staining of samples incubated with Fab IV.3, using a secondary RαM-FITC antibody, reveal that the marker is present on the majority of cells (E).

### The interaction network of CD13, Syk and CR3 (CD11b/CD18) functional partners contains 76 proteins

The previous results showed that crosslinking CD13 on MDMs induced the high affinity conformation of CD11b. Next, we turned to bioinformatic databases to assemble an interaction network comprised of functional partners of CD13, CR3 and Syk, a key signalling kinase in the immune system, particularly in myeloid cells, in order to propose a sequential mechanistic model for the inside-out signalling pathway that could account for the activation of CR3 following CD13 crosslinking.

To determine the potential set of proteins and pathways that participate in the CD13-CR3 inside-out-signalling cascade we constructed an interaction network using information from public databases, literature, and previous experimental work from our laboratory. Given the high number of potential candidates, network nodes were selected by predicted interaction score, biological function, and presence in the target cell type.

A functional protein interaction network of CD13, Syk and CR3 was assembled selecting the proteins with the highest combined scores (0.8 or more) from the STRING database (*32*), as well as previously determined experimental interactions. Data mining STRING element, and the databases GeneCards (*33*) and PubMed (*34*) were used to confirm that the chosen proteins were present in the myelomonocytic lineage. Fig. S1 presents the main ontology clusters for the selected proteins. For those interrogation nodes that resulted in more than 50 proteins with combined scores ≥0.8, the top 50 molecules were analysed.

Using Syk as the interrogation query we obtained a first layer of interactions of 158 proteins with STRING combined scores above 0.9. Twenty-nine entries were selected according to the established criteria, i.e., representing CD13 and/or CR3 known functional interactors or potential elements for the inside-out signalling pathway connecting the two of them. Two of these proteins were also selected in the CD11b and CD18 analyses. Fig. S2 includes the proteins selected to assemble the network, and the Venn diagram allows identification of those molecules common to two or more interrogation queries. In the case of Syk, two of its interactors were also common with CD11b and CD18.

Polypeptide chains forming CR3 (CD11b (ITGAM) and CD18 (ITGB2)), were also subjected to this type of analysis. For CD11b, the first layer of interactions with STRING combined scores above 0.9 consisted of 167 proteins, resulting in 22 molecules of interest. Ten of these were also selected for CD18, as well as the two previously mentioned for both Syk and CD18.

Using CD18 as the interrogation query resulted in 184 interactors with a combined score ≥0.9. Twenty-five molecules of interest were chosen, 13 of which were exclusive to CD18, and the rest were shared with Syk and CD11b, as aforementioned.

Using CD13 as the interrogation query yielded 27 molecules with STRING combined interaction scores of 0.8 and above, these were filtered to 4 proteins of interest using the criteria of being either downstream signal inhibitors or enhancers, adhesion molecules or co-receptors which, following text mining, might provide information on the signalling pathways necessary for interaction with CD13. Finally, 12 proteins for which interaction with CD13 was previously experimentally determined (SYK, GRB2, PI3K, FAK, IQGAP1, SRC, JNK, p38, MEK-1, PKC, ERK 1/2 and SOS1) were added to the molecules of interest (*5, 35, 36*).

Fig. 3 depicts the interaction network obtained, consisting of 76 non-redundant proteins. Of note, pink lines and bubbles represent interactions experimentally determined, including the ones contributed by this study. A detailed list of all proteins in the network, their main characteristics, and their corresponding interrogation nodes, is presented in Table S1.

**Fig. 3.**
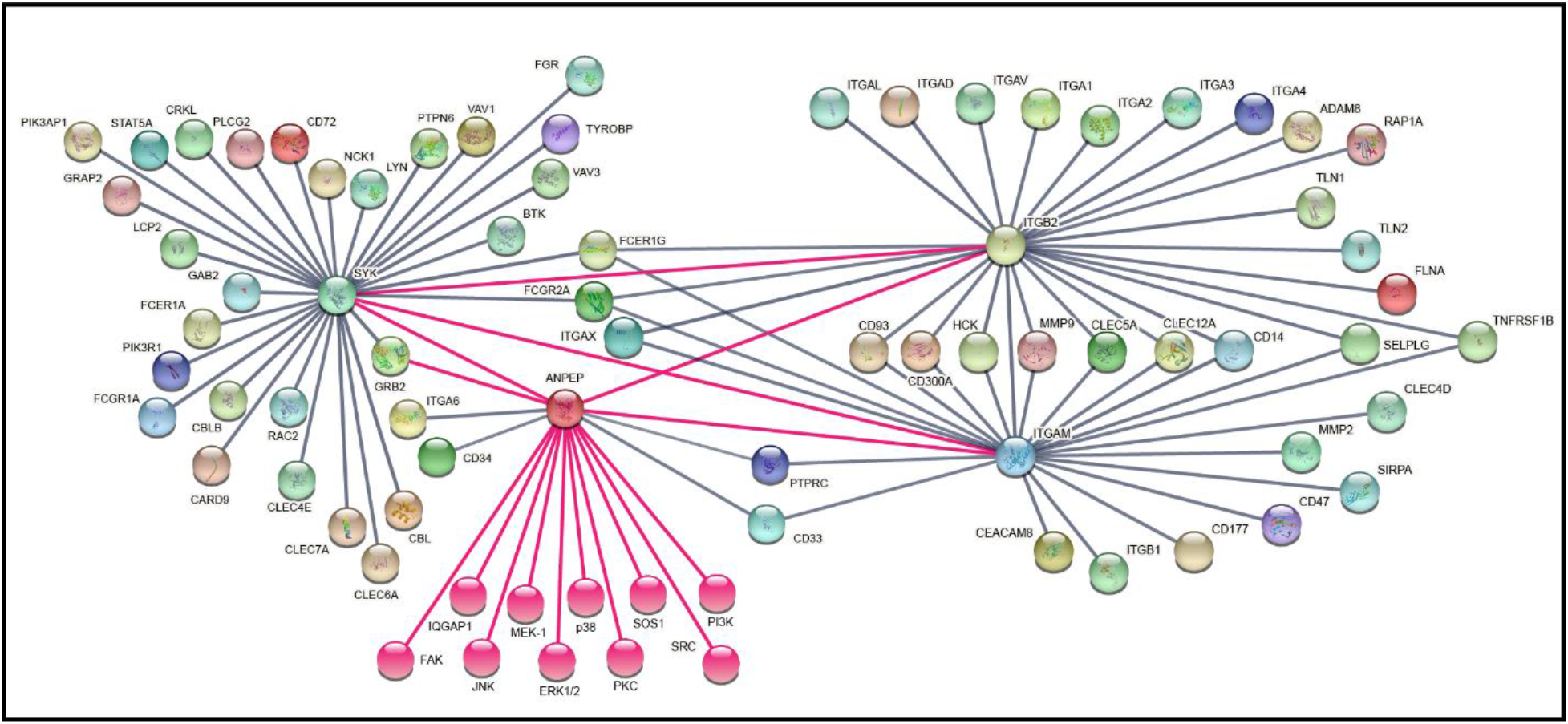
The interaction network of CD13, Syk and CR3 (CD11b/CD18) functional partners comprises 76 proteins. For Syk, the first layer of interactions with a score above 0.9 consisted of 158 proteins, similarly filtered to 29. Both CR3 polypeptide chains were analysed. 167 and 184 proteins with a score above 0.9 were found for CD11b and CD18, respectively. From these, 10 were exclusive to CD11b, 13 were exclusive to CD18, 10 were shared between the two of them and two with Syk as well. As for CD13, the first layer of interactions consisted of 27 proteins, which were narrowed down to four. Pink lines and bubbles represent the interactions experimentally determined in our laboratory and others, which adds 12 more to the proteins of interest. Non-redundant results are depicted.

Next, based on our interaction network, we constructed a sequential mechanistic model of the CD13-CR3 inside-out signalling pathway.

### Sequential mechanistic model of the CD13 to CR3 (CD11b/CD18) inside-out signalling pathway

Unlike many other adhesion and phagocytic receptors, CD13 has only a short cytoplasmic tail with no canonical signalling motifs (*36*). However, this is not an impediment for acting as a trigger of phosphorylation cascades. Upon antibody crosslinking, CD13 dimers are phosphorylated on their intracellular portion (Tyr6), most likely by Src (*5*). Therefore, after demonstrating that CD13 crosslinking activated the integrin CR3, we blended the insights provided by these experiments with the information from our CD13-Syk-CR3 interaction network to propose a sequential mechanistic model for the CD13-CR3 inside-out signalling pathway, a depiction of which can be seen in Fig. 4.

**Fig. 4.**
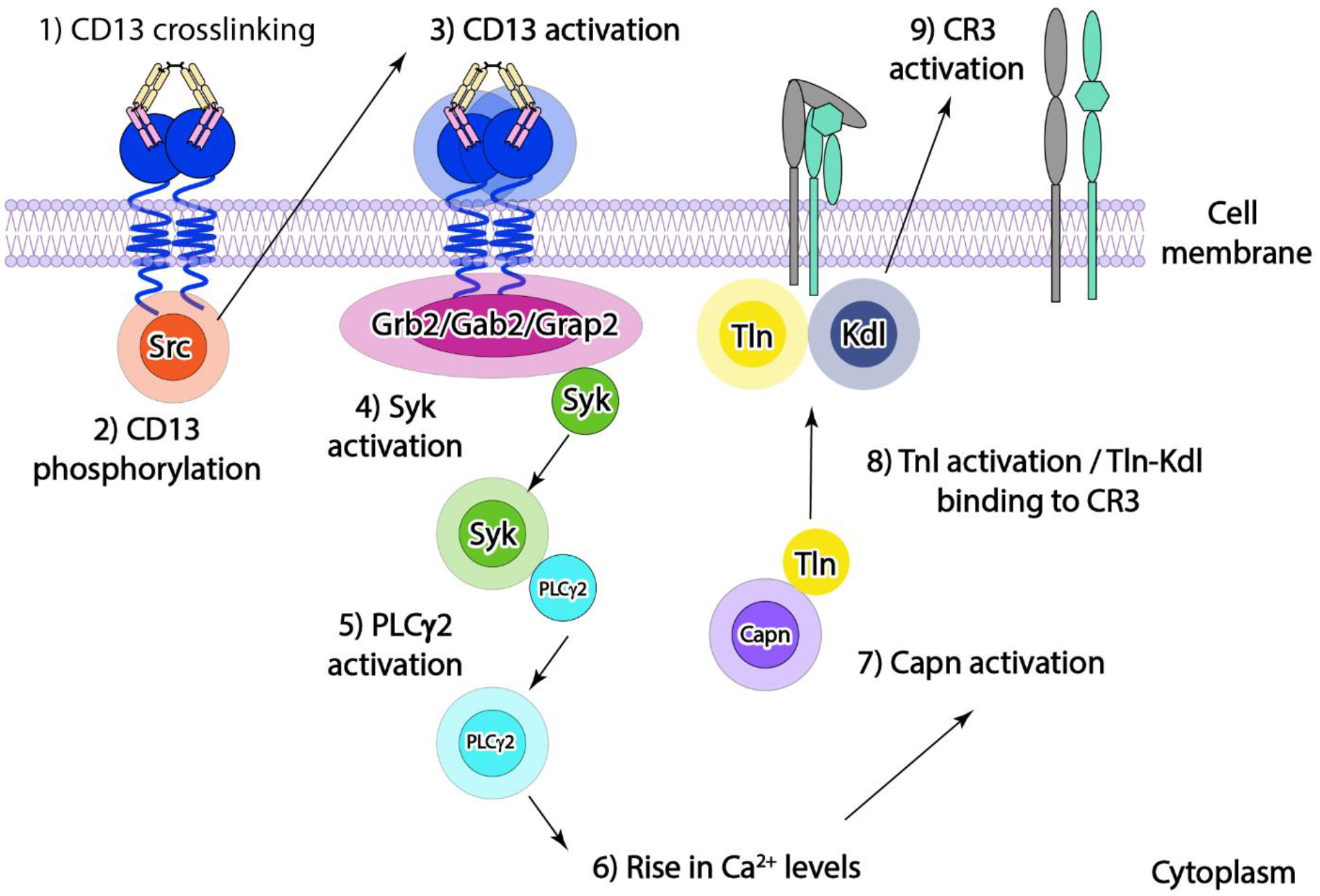
CD13 to CR3 inside-out signalling pathway. Experimental and bioinformatic data were used to propose an initial CD13-CR3 inside-out signalling pathway. This schematic representation depicts the main events of this signalling cascade. In it, CD13 is activated upon crosslinking and Src-mediated phosphorylation, next, an adaptor protein binds this receptor, allowing Syk to autoactivate. Syk phosphorylates PLCγ2, inducing the PI3K pathway and, consequently, a rise in Ca^2+^ levels. This activates the peptidase Calpain, which cleaves Talin, then permitting its interaction with Kindlin and subsequent CR3 activation.

Tyr6 phosphorylation on both crosslinked CD13 molecules may create a docking site for scaffolding or adaptor proteins like Grb2, Grap2 or Gab2. Subsequently, other molecules like Sos1, Syk and other non-receptor tyrosine-kinases and phosphatases such as SHP-1 (PTPN6) could be recruited. The next step would be the auto-activation of Syk and detachment from the adaptor protein, although it is also plausible that Syk auto-phosphorylates upon binding to CD13 directly on the Tyr6 docking site, as it does on ITAMs during FcγRs signalling. Syk may activate PLCγ2, which produces IP3 and DAG. As a consequence of IP3 production, Ca^2+^ is released from the endoplasmic reticulum. One of the many Ca^2+^-dependent enzymes is Calpain, which then would cleave and activate Talin (TLN1/2) (reviewed in (*37*)). Then, Kindlin and Talin destabilize the CD18-CD11b interaction, specifically, because activated Talin interferes with the salt bridge between the integrin subunits (*38*). This would lead to CR3 transitioning from the low-affinity to the high-affinity state.

We decided to evaluate this hypothetic signalling pathway by experimentally inhibiting a key protein in many myelomonocytic cascades: Syk. If our assumptions were correct, the activation of CR3 would be diminished, if not completely abolished, by incubating the MDMs with BAY, a Syk inhibitor.

### Syk inhibition resulted in a higher number of CR3 (CD11b/CD18)-activated cells

In order to evaluate the hypothesis of Syk as partly responsible for the activation of CD11b following CD13 crosslinking, MDMs were first incubated in serum-free medium for different time periods with BAY, before assessing CD13-induced activation of CR3.

The cells analysed in the flow cytometer were gated as described in figures one and two. Fig. 5A shows that histograms from cells incubated with BAY and, without antibodies or only with the secondary antibody used for crosslinking and stained with anti-CD11b(activated)-FITC, were not displaced in comparison with the unstained control, meaning that there was no nonspecific CD11b activation generated by either handling the cells or the secondary antibody alone. However, when MDMs were incubated with Fab fragments of a primary anti-CD13 antibody (Fab C) and crosslinked with F(ab)’2 GαM fragments, a clear signal was detected as depicted in the representative histograms in Fig. 5B, where CD11b was activated in average in 30% (±16.7) of the control cells population. In contrast to what was expected, when cells were incubated with BAY, this percentage rose to an average of 54% (±14.5). Fig. 5C is the graphical representation of all tested pairs of CD13-crosslinked cell samples in the presence of BAY and their corresponding controls. Median fluorescence intensity (MFI) also rose significatively in the presence of BAY, from an average of 487(±78) to 574 (±115) arbitrary units (a.u.) (Fig. 5D). The percentage of CD11b activation and MFI were analysed using paired two-tailed t tests, which confirmed the statistical significance of the differences.

**Fig. 5.**
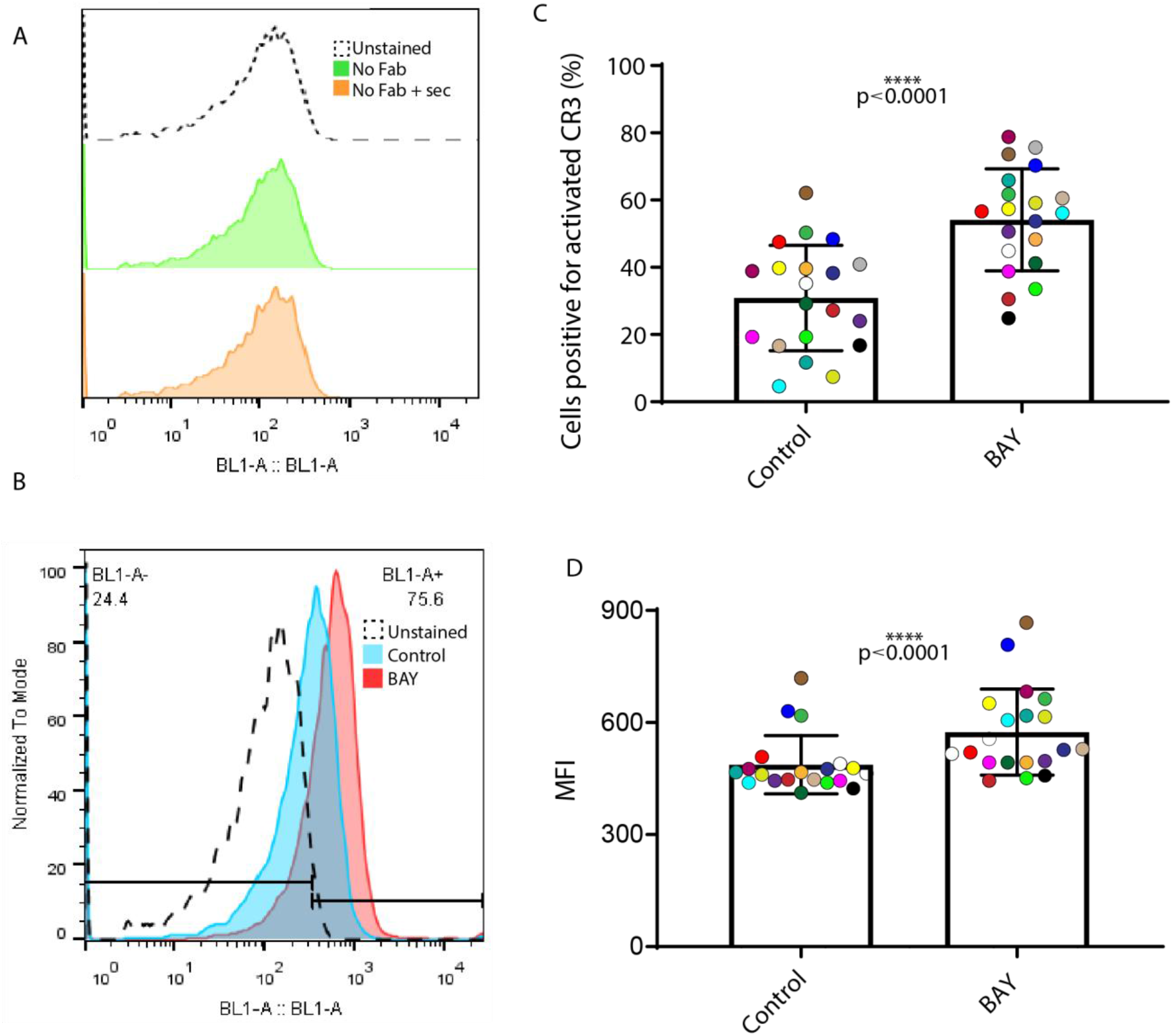
The Syk inhibitor, BAY, elevates the percentage of CR3-activated cells in CD13- crosslinked MDMs. Cells were first gated for size and granularity, then for singlets and finally for median fluorescence intensity in the BL1 (FITC) channel (A and B). BAY-incubated controls are shown in panel A. In panel B is possible to appreciate the difference in CR3 activation in representative histograms from a 0h sample and its corresponding control. Panel C shows the percentage of CR3 activation cells plotted for all 20 pairs of matching samples and controls, with each pair colour-matched. An average of 30.8% (±16.7) of control cells showed CR3 activation after CD13 crosslinking; this number rises to 54.1% (±14.5) of the population on average, when cells are incubated with BAY. Panel D shows MFI from the same 20 pairs of samples equally colour-matched. Average MFI from controls was 487(±78) a.u., increasing to 574 (±115) a.u. in cells incubated with BAY. Two-tailed paired t-test confirmed that the differences were statistically significant, p<0.0001. CI 95% was 15.4 to 31 for percentages and 55.90 to 118.1 for MFI.

These observations suggested that our decision to evaluate the role of Syk, a key molecule for the immune system, especially in the myelomonocytic lineage, was appropriate since its inhibition revealed the inhibitory loop of the CD13 to CR3 inside-out signalling pathway. In order to adjust the sequential mechanistic model to one in which Syk is a signal controller, we revisited our interaction network to include at least one element that, upon phosphorylation, restrains the activation of CR3.

### Syk is an inhibitor of the CD13-CR3 (CD11b/CD18) inside-out signalling pathway

Given this new experimental information, we modified the sequential mechanistic model initially proposed to include the suggested Syk-mediated inhibitory loop. The improved model included 12 proteins and 11 interactions (Fig. 6). It integrated multiple mechanisms like the phosphorylation of the ubiquitin ligase Cbl and the subsequent ubiquitination of one or more proteins from the signalling cascade. These results highlighted the importance and pragmatism of combining experimental and bioinformatic approaches to efficiently decipher the complex inhibitory or enhancing mechanisms that underlie signalling pathways.

**Fig. 6.**
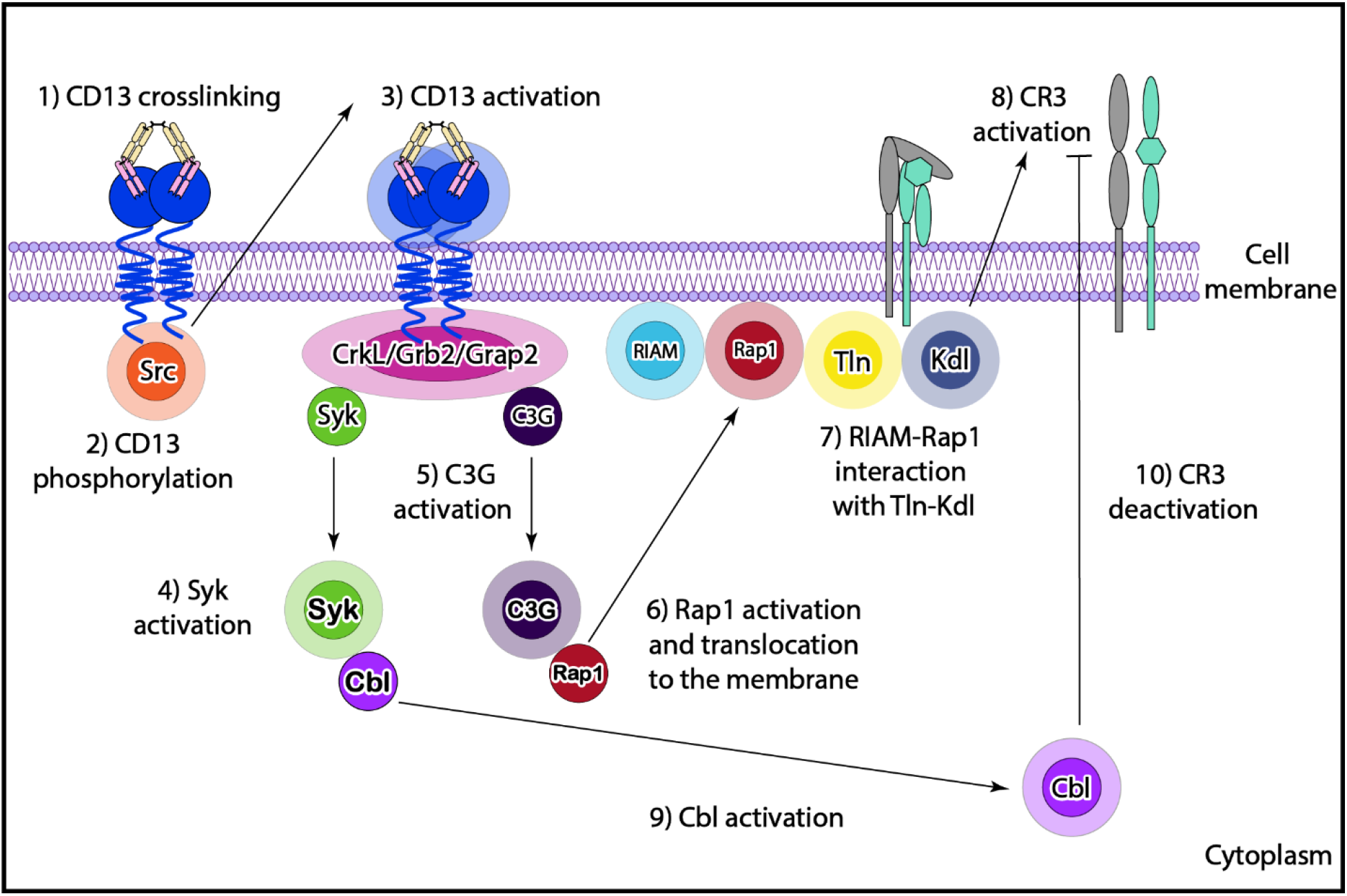
Syk is an attenuator of the CD13-CR3 inside-out signalling pathway. Experimental data from CR3-activation experiments in the context of Syk inhibition, suggest that this molecule is a regulator of the CD13-CR3 pathway, rather than an activator of it, as initially thought. The new hypothesis preserves the initial steps from the one originally proposed: first, CD13 activation ensues after antibody crosslinking and Src-mediated phosphorylation; secondly, adaptor molecule docking on CD13’s cytoplasmic tail, followed by Syk auto-activation. Then, Ras GEF C3G is activated though its interaction with and adaptor molecule, allowing Rap1 activation. The RIAM-Rap1-Talin-Kindlin axis then assembles, producing CR3 activation. Finally, it is conceivable that Syk activates a member from the ubiquitin ligase Cbl family, which deactivates CR3 by ubiquitination of one or more pathway components.

Experimental data from cells incubated with BAY showed that Syk inhibition resulted in a higher number of cells with an activated-CD11b positive signal. Thus, the activation of CR3 must proceed through a different mechanism than the one initially hypothesized.

Nevertheless, it was possible to retain the suggested initial events following CD13 crosslinking, that is, Tyr6 phosphorylation on both CD13 molecules by Src, creating a docking site for adaptor molecules like Grb2, Grap2 or CrkL. Then, Syk self-activation either prior to adaptor protein- CD13 interaction, via auto-phosphorylation by binding the CD13 phospho-tyrosines as it does with ITAMs on Fc receptors (reviewed in (*39*), or by binding the adaptor molecule at its SH2 or SH3 domains (*40*), after such molecules assemble in a CD13-adaptor complex.

In parallel, the Rap GEF, C3G would activate via its interaction with the adaptor protein, most likely CrkL (*41*), then it may catalyse the GDP/GTP exchange in Rap1. After phosphorylation by Fak and Src, the protein RIAM would bind to Rap1 through its Ras association domain, and to PIP2 within the cell membrane via its Pleckstrin domain (*42*). Alongside its binding to RIAM, Rap1 also anchors to the membrane, and interacts with the Talin-Kindlin axis, which would drive the transition of CR3 from its inactive to its active state. Rap1 is a key node in the inside-out activation of phagocytic integrins like CR3. This small GTPase is where the signals coming from different receptors such as those for some chemokines and cytokines, and certain TLRs, converge and result in integrin activation (*43*).

Finally, unlike in the originally proposed mechanism, Syk would phosphorylate members of the ubiquitin-protein ligase Cbl family, namely Cbl and Cbl-b, present in our interaction network (*43*). These ligases may ubiquitinate one or more components of the signalling pathway, including CD13. This would induce an inhibitory loop that halts the signal transduction inducing CR3 activation. Syk inhibition potentially interferes with this mechanism resulting in a higher degree of CR3 activation, as seen in our experiments with MDMs incubated with BAY.

## DISCUSSION

Over the years the description of CD13 has gone from a marker of leukaemia to a co-receptor, to a moonlighting enzyme in its own right. This work contributes to show that CD13 is able to elicit not only outside-in signalling, but to triggered at least one inside-out signalling pathway, thus activating another immune receptor CR3 (CD11b/CD18) in a specific manner. Our data are in line with similar observations such as those from Ortiz-Stern and Rosales (*44*), who report the activation of a β1 integrin after CD32b (FcγRIIb) crosslinking. This demonstrates once more that, even in the absence of canonical signalling motifs, CD13 induces cell phenomena comparable to classical phagocytic receptors.

We assembled a functional protein interaction network for CD13, Syk and CR3 using previously published experimental data from our lab and others, as well as bioinformatic resources like STRING, GeneCards and PubMed databases. This network comprised 76 proteins and allowed us to filter the potential elements for the CD13-CR3 inside-out signalling pathway. The network was used to build a model for the sequence of events that concatenate CD13 crosslinking and CR3 activation. The hypothetical pathway was experimentally tested, and the model further refined, swiftly expanding the mechanistic understanding of a complex phenomenon such as a inside-out signalling cascade. This highlights the synergy between in silico and experimental approaches. A summary of our workflow can be seen in Fig. S3.

Each step in our sequential mechanistic model (Fig. 4) and its subsequent modification (Fig. 6), was supported not only by STRING predicted interactions with the interrogation queries, but also by previous experimental data. This made both pathways theoretically conceivable. For example, we initially chose Grb2 as an adaptor molecule bridging CD13 and Syk because our group found in previous studies that crosslinked CD13 co-precipitates with Grb2, and this molecule associates with Shc, Src, Syk and SHP-1 during inside-out signalling between CD32a and αIIb/β3 integrin in human platelets (*25, 45*). We also suggested the activation of the recognized Syk substrate PLCγ2, whose activity ultimately sparks the release of Ca^2+^ from the endoplasmic reticulum (*39*). Of note, Ca^2+^ release is a known effect of CD13 crosslinking by specific antibodies (*4*). Additionally, similar inside-out signalling pathways are induced when PSGL-1 from human neutrophils binds its endothelial ligands, P- and E-selectins, resulting in the activation of β2 integrins CR3 and LFA-1 (CD11a/CD18) (reviewed in (*9*)).

The feasibility of this proposal was also supported by the findings of Zheng et al.(*46*), who reported that stimulation of glycoprotein VI leads to the activation of αIIBβ3 integrin in platelets, an effect dependent on the Syk-mediated phosphorylation of PLCγ2. Moreover, the activation of αIIBβ3 integrin is prevented by BAY 61-3606, the same chemical inhibitor used in our experiments. testing of our model, we established that the non-receptor tyrosine kinase Syk is an attenuator of the CD13-CR3 pathway, as its inhibitor BAY causes a statistically significant increase in both the number of cells bearing activated CR3, and the number of activated CR3 molecules on their surface. Thus, our initial model was modified accordingly. Among the adjustments to the model were: i) the inclusion of the adaptor molecule CrkL, present in our interaction network, as a bridge between p-CD13 and the rest of the cascade, ii) the participation of the Rap GEF, C3G as the GDP/GTP exchanger for Rap1, justified by the presence in our interaction network of both its activator and substrate, CrkL and Rap1, respectively and, iii) the suggested ubiquitination of components of the cascade by a member of the Cbl family, which are Syk-stimulated ubiquitin-ligases. Of note, such chemical modification does not necessarily mean that targeted proteins would be degraded, they could be deubiquitinated in late endosomes and recycled back to the cytoplasm or cell membranes, as it occurs with a proportion of EGFR after signalling termination (reviewed in (*46*)).

Hence, when Syk was inhibited by BAY, this attenuating mechanism could not proceed in a timely manner, and further CR3 activation was registered compared to the samples incubated without BAY. This possibility is not unreasonable since Cbl-deficient murine bone marrow derived mononuclear phagocytes display enhanced β2 integrin-mediated adhesion during inside-out dependent activation (*47*).

The fact that more than one pathway was proposed for CD13-dependent CD11b activation demonstrated that the number of candidates in the interaction network was rich enough to supply an ample number of protein options for different scenarios, provided that each individual step of the proposed pathway had been previously reported, as in the case of our mechanistic model. It also showed that the outcomes from different stimulation conditions are finely-tuned. In the case of CD13, we see two different roles for Syk, depending on the nutritional stress of the cell. For example, in our experiments when CD13 was crosslinked with antibodies after a period of starvation (serum-free incubation), Syk functions as part of the inhibitory mechanism of the CD13-CR3 pathway. However, in previously reported data, when cells are incubated in serum-supplemented medium, along with a phagocytic prey, phagocytosis, and reactive oxygen species (ROS) production require that Syk acts as an activator (*3*).

This makes sense from a bioenergetic and evolutionary point of view, since the cell orchestrates its responses using almost the same group of proteins instead of producing an entirely different set for each function, i.e. those in which Syk acts as an activator (like phagocytosis and ROS production), and those in which Syk is an inhibitor (like CR3 activation). Furthermore, the link between CD13 and CR3 is also supported by *in vivo* evidence, as both molecules can be found together in functional microdomains within the cell membrane called lipid rafts (*29*). These structures are key to cell signalling since they bring components of specific pathways close together, thus, decreasing the possibility of fortuitous activation or blocking of signals from other cascades (reviewed in (*48, 49*)).

Our results also exposed a further level of complexity for the CD13-CR3 pathway as CD13 crosslinking activated CR3 in a fraction of the treated MDMs population, and the inhibition of Syk augmented this fraction. However, such a rise was not observed across the entire population in any of the tested conditions. There are multiple possible explanations for this observation. The first one is that, the overlapping activity of other kinases, conceivably Src-family kinases like Lyn, may substitute for Syk when it is inhibited, by phosphorylating the suggested target in this pathway, Cbl (Cbl/Cbl-b) (*50*). A second possibility is that, as the antibody used for the assessment of CD11b activation (commercial mAb CBRM1/5) did not recognize its intermediate affinity state, such a change was not quantified even though it may also be a consequence of triggering the CD13-CR3 signalling pathway. This is supported by the findings of Chung and colleagues (*51*), who detect both high- and intermediate-affinity specific epitopes on β2 integrins upon TLRs stimulation in THP-1 monocytic cells using different monoclonal antibodies. A third possibility is that, in the absence of an additional stimulus, the CD13-induced activation signal was not sufficient to activate CR3 in every cell, as occurs in transformed human brain microvascular endothelial cells. When these cells are exposed to IL-1β, activation of α5β1 integrin rises, binding to fibronectin is enhanced and the subsequent signalling is increased, compared to unexposed cells (*52*).

Our results suggest that CD13 and CR3 bring about one or more cellular functions as an ensemble. One of them is adhesion. CD13 has long been implicated in pro-adhesive events for example, it induces homotypic aggregation in myeloid cells (*53, 54*) and plays a role in the invasiveness of osteosarcoma cells in vitro and in vivo (*55*). The adhesive properties of CR3 (CD11b/CD18) are well known, as is its tendency to be activated after the stimulation of other receptors (inside-out signalling), in a variety of settings both beneficial and detrimental to the host. For example, CD44-mediated phagocytosis in murine macrophages triggers and is partially dependent on CD11b activation (*56*), potentially resulting in the destruction of the phagocytic prey. In contrast, recognition of human neutrophil antigen 3a by auto-antibodies triggers CD11b activation, causing neutrophil accumulation in the pulmonary microvasculature of some blood transfusion recipients, driving severe transfusion-related acute lung injury (*57*).

It is therefore conceivable that CD13 and CR3 take part in the same adhesion chain of events during inflammation-related transendothelial migration; our group previously reported that CD13-mediated adhesion to endothelial cells is integrin-independent (*36*), although it is important to note that CD11b is not among the integrins evaluated. Thus, it is possible that CD11b indeed contributes to this process, but at this stage it cannot be confirmed or ruled out. This would partially explain the observation that upon in vivo CD13-ligation transendothelial migration is impaired (*36*). Given that, as we demonstrated, CD11b activation is a consequence of CD13 stimulation, persistent CD13 engagement would render active CR3 in constant contact with its endothelial ligands like ICAM-1 and -2 (*11*), maintaining the cell in arrest in a Kindlin-dependent manner (*12*), or like with JAM-A, JAM-C and RAGE, triggering outside-in signalling and causing polarization and spreading (*58*) albeit to an extent that would block extravasation.

Another function possibly coordinated between CD13 and CR3 is phagocytosis, as both perform this cellular function. Licona-Limón and colleagues (*3*) demonstrated that CD13 is a primary phagocytic receptor, as phagocytic prey selectively directed towards CD13 are engulfed by human macrophages and THP-1 monocytes at the same rate as they engulf FcγR-directed particles. Furthermore, when CD13 is expressed on HEK293 cells, which normally do not express this receptor, they are able to internalize the same type of phagocytic prey. As for CR3, it is well known for its participation in complement-mediated phagocytosis. The complement system is a series of circulating proteins mainly synthesized in the liver. The three complement activation pathways (classical, lectin and alternative) can be summarized as a series of successive protein cleavages resulting in the production of a C3 convertase, which splits the C3 protein into C3a and C3b fragments. C3b opsonizes pathogens and is further processed giving rise to iC3b and C3dg (reviewed in (*10*)). iC3b-opsonized pathogens are recognized and phagocytosed through CR3. CR3-mediated phagocytosis is synergistically enhanced by other receptors, for example, *Francisella tularensis* is internalized in concert by CR3 and CR1, each binding their own ligands on the surface of the bacterium (*59*), or *Borrelia burgdorferi*, whereby internalization by human macrophages is orchestrated by CR3, CD14 and scavenger receptors (*60*). As CD13 also acts as co-receptor to other phagocytic receptors like FcγRs and Mannose receptors (*26, 61*), this may also be the case for its functional interaction with CR3.

One limitation of this study is that the functional consequences of CR3 transitioning from a low-affinity to a high-affinity state in response to CD13 stimulation were not explored. Therefore, we can only speculate about the possible scenarios where CD13-mediated CR3 activation could be biologically relevant. A second limitation is that, given the number of nodes in the protein interaction network, it may have been viable to propose one or more extra sequential mechanistic models for the CD13-CR3 signalling pathway. Such models could be a convenient backup resource in case that, upon further testing, the final model presented in this article proves not entirely accurate.

As detailed out by Santos et al. (*35*) and reviewed by our group (*62*) CD13 is part of a group of ectopeptidases, along with CD157, CD73, CD38 and CD26, that initiate signalling events upon stimulation. Despite the need for extra accessory proteins, the existence of receptors without tyrosine-kinase activity (nRTKs) like these ectopeptidases, may have been retained during evolution, providing a tighter cell activation control than receptor tyrosine kinases (RTKs). nRTKs are not prone to auto-phosphorylation upon stochastic encounters in the cell membrane.

In contrast, spontaneous activation is possible with RTKs, which is a great disadvantage in conditions where they are overexpressed as it can result in disease development. For example, human epidermal growth factor receptor 2 (HER2) is a RTK and its overexpression is linked to ovarian, prostatic, gastric, lung and breast cancers (*63*). Moreover, activation of HER2 is a recognized mechanism of resistance to endocrine treatment in several experimental models (*64*).

The results of our study highlight the fine-tuning of both inside-out and outside-in signalling involving a single receptor, CD13, as many of the same proteins participate in either pathway, however, with quite different consequences. Syk is a key activator in the outside-in signalling following CD13 and CR3 engagement (*3, 16*), however, we demonstrated the inhibitory role that this same soluble tyrosine kinase has in the inside-out communication between both receptors.

CD13 is overexpressed in many cancers, whereby adhesion and cell motility, a mechanistically closely related phenomenon, contribute decisively to tumour progression (*65–67*). Therefore, future research should be directed towards the functional impact of CD13 crosslinking on CR3- mediated adhesion and phagocytosis, i.e., the identification of the functions mediated in concert by these receptors. It will also be necessary to assess the participation of other components of the CD13-CR3 signalling pathway in vitro and, eventually in vivo. One option is to use a CRISPR- Cas9 screening strategy, in which each of the genes coding for the nodes can be interrupted in individual immortalized cells, and then expanding these into cell lines. This would make it possible to ascertain characteristics of the signalling pathway such as event timing or the consequences of the absence of each protein. For example, according to our proposed sequence of events, the first protein to be tested should be Src, which can be initially done by chemically inhibit it and observe whether the activation of CR3 is affected, if so, then a genetic strategy can be applied. A similar approach would be useful to determine the kinases whose activity overlaps with that of Syk, in which case a cell culture where Syk is knocked down would be convenient to test specific inhibitors for different Src family kinases.

Additionally, the phosphorylation at serine 8 and 10 in the cytoplasmic tail of CD13, which has not been reported yet, should be evaluated as it could add extra docking sites for accessory proteins. Such specifics could provide the basis for the design of therapies that inhibit or enhance cellular activities to prevent spread of cancers in which CD13 is overexpressed.

Finally, we wish to stress the convenience of this interdisciplinary approach for signalling transduction studies. In our experience, bioinformatic tools were fundamental for broadening the reach of a few carefully designed key experiments, and to delineate not one, but two possible signalling cascades based on them. This rationale has the potential to provide researchers with a pipeline that has proven to save time and resources in our hands, as well as help refine concrete future goals.

## MATERIALS AND METHODS

### Reagents and antibodies

RPMI-1640 medium was purchased from Gibco Life Technologies (Carlsbad, CA, USA). Recombinant human (rh) M-CSF was from PeproTech (Cranbury, NJ). Lymphoprep was from Axis-Shield PoC AS (Oslo, Norway). All culture media were supplemented with 10% heat inactivated FBS (Invitrogen, Carlsbad, CA, USA) unless otherwise stated, 2 mM L-glutamine, 100 μg/ml streptomycin, 100U/mL penicillin (Sigma-Aldrich, St. Louis, MO, USA), 1 mM sodium pyruvate solution and 1% MEM non-essential amino acids solution(100X) (Gibco by Life Technologies, NY, USA). Murine monoclonal IgG1 anti-human CD13 (Mab C) and anti-human CD32 (Mab IV.3) were produced and purified in our laboratory from supernatants of the corresponding hybridomas (*4*). Fab fragments were prepared from the purified antibodies with immobilized Ficin (Pierce, Rockford, IL), following the manufacturer’s instructions. Murine monoclonal FITC anti-human CD11b (activated) antibody (IgG1, clone CBRM1/5) was from Biolegend (San Diego, CA). Goat anti-mouse (GαM) polyclonal IgG F(ab)’2 fragments were from Jackson ImmunoResearch (West Grove, PA). Polyclonal FITC rabbit anti-mouse antibody was purchased from Thermo Fisher Scientific (Waltham, MA). BAY 61-3606 was from Sigma-Aldrich (St. Louis, MO, USA).

### Cell Culture

All experiments carried out with cells from human donors were performed following the Ethical Guidelines of the Instituto de Investigaciones Biomédicas, UNAM, Mexico City, Mexico.

Human peripheral blood mononuclear cells (PBMCs) were isolated from anonymous healthy male donors’ buffy coats obtained from the blood bank at Instituto Nacional de Ciencias Médicas y Nutrición Salvador Zubirán, Secretaría de Salud, Mexico City, Mexico by gradient centrifugation with Lymphoprep as previously described (*4*). For monocyte isolation, PBMCs were washed three times with PBS, pH 7.4 by centrifugation at 400 g for 10 min. After the last wash, cells were resuspended in serum-free RPMI-1640 medium complemented as described before and were seeded (5–6 × 10^7^ PBMCs/plate) in 100 mm × 20 mm cell culture-treated polystyrene culture dishes (Corning, New York, NY, USA). Cultures were maintained in a humidified atmosphere at 37°C with 5% CO2 for 1 h, to allow monocytes to adhere to the plastic plate. Non-adherent cells were eliminated by gentle washing, and adherent cells, enriched for monocytes (≥95% purity, as determined by flow cytometry using CD14 as a marker of the monocytic population, data not shown), were cultured for 7-10 days for differentiation into macrophages, in RPMI-1640 medium with 5 ng/ml rh M-CSF complemented as described before, at 37°C. For experiments, macrophages were harvested by firm and gentle cell scraping.

### CR3 activation

MDMs (5×10^5^) were incubated in 6-well plates in serum-free supplemented RPMI-1640 medium for a maximum of 12 h with or without 10 μM BAY, a highly selective and widely used Syk inhibitor (*68–70*). One well was harvested by gentle cell scraping at 0, 3, 6 and 12h. From each harvested sample 0.25×10^6^ MDMs were incubated in 0.2 ml serum-free supplemented RPMI- 1640 medium with 2.5 μg of mAb C (anti-CD13) Fab fragments for 30 min at 4°C. Cells were washed three times with the fresh medium and incubated with 4 μg of GαM F(ab)’2 fragments for 30 min at 4°C. Immediately after, cells were incubated for 10 min at 37°C. Finally, cells were fixed with 1% paraformaldehyde (PFA) for 10 min at RT.

### Flow cytometry

To quantitate CR3 activation, fixed samples were washed two times with cold PBS and stained with 20 μl of a 1:20 dilution of murine monoclonal FITC anti-human CD11b (activated) antibody (IgG1, CBRM1/5) for 40 min at 4°C. Cells were washed three times with cold PBS.

Staining for CD13 or CD32 (FcγRII) was performed on MDMs by incubation in 10 μM anti- CD13 Fab C or anti-CD32 Fab IV.3 in serum-free supplemented RPMI-1640 medium for 30 min at 4°C. Cells were washed three times with the same medium and incubated with 1:400 GαM- FITC antibody for 30 min at 4°C, then washed three times with cold PBS and fixed with 1% PFA for 10 min at RT. Fluorescence intensity was measured by flow cytometry (Blue/violet Attune cytometer, Applied Biosystems-Thermo Fisher, Waltham, MA). Flow cytometry data are displayed as percentages of gated (ssc vs fsc, and singlets) positive cells compared to non-treated controls

### Theoretical cell signalling interaction network assembly

We constructed the functional protein interaction network of CD13, Syk and CR3 and their closest partners using combined interaction scores from STRING (*32*). A functional association in this context means either physical contact, participation in the same metabolic pathway and/or cellular process (*71*). STRING scores are indicators of the likelihood of an interaction, given currently available evidence in the database, which include gene neighbourhood, gene fusions, gene co-occurrence, experimental evidence, curated databases, text mining, and protein homology. Each type of evidence gives rise to an individual score for each pair of proteins.

STRING computes combined scores by integrating the individual scores and correcting for the probability of randomly observing the interaction. Scores rank from 0 to 1, with 1 being the highest possible result.

The search for functional partners was done individually for each interrogation query (CD13, CD11b, CD18 and Syk) and focused on human proteins. High confidence scoring molecules (0.8 and above) from the first layer of interactions with the query were considered. The resultant group of proteins were filtered based on the requirements for this particular inside-out signalling pathway: non-receptor kinases, adaptor proteins able to bridge CD13 to other components of the pathway, specially Syk, and inhibitory molecules, like protein phosphatases or ubiquitin ligases. In some cases, other interacting receptors were considered, as they may provide insight into the reported mechanisms for this type of interaction. Namely those similar to the studied receptors, CD13 and CR3: metalloproteases, phagocytic receptors, integrins and other adhesion molecules. To ensure the quality and specificity of the network text mining STRING element, The GeneCards website (*33*) and the repository PubMed (*34*) were used to ascertain the suitability of each selected protein, i.e. to confirm the function of each node, as well as its gene and protein expression in myelomonocytic cells. Finally, the interaction network was manually curated according to experimental evidence gathered from previous publications.

### Statistical analysis

For receptor crosslinking experiments in the absence of BAY, statistical analysis was performed by means of one-way ANOVA followed by Dunnett’s multiple comparisons test or a paired two-tailed t test in the case of BAY-incubated cells and their controls. P values below 0.05 were considered significant.

## Supplementary Materials

Fig. S1. Molecular ontology within the protein interaction network

Fig. S2. 65 non-redundant proteins were deemed of interest after the interrogation of databases using CD13, Syk and CR3 (CD11b/CD18) as queries

Fig. S3. Workflow

Table S1. Description of the 76 proteins contained in the CD13, CR3 (CD11b/CD18) and Syk interactions network

## Acknowledgments

The authors thank the technical support from Dr. Claudia A. Garay-Canales, PhD, with cell biology procedures and from Carlos Castellanos-Barba, MSc, at the National Laboratory of Flow Cytometry (LabNalCit), Instituto de Investigaciones Biomédicas, UNAM. The authors are also deeply grateful to Drs. G. Erandi Pérez-Figueroa, Marco A. Alfonzo-Mendez and Bruce Alberts for their generous suggestions to improve the quality of the manuscript. Laura Díaz-Alvarez is a doctoral candidate in the Posgrado en Ciencias Biológicas UNAM program, this article constitutes a requirement for the attainment of her PhD degree.

## Author contributions

Conceptualization, methodology: LDA, MEMS, EO

Data curation, formal analysis, investigation and writing of the original draft: LDA Project administration: LDA, EO

Supervision: EO, MEMS

Visualization, resources, validation and review/editing of manuscript: LDA, MEMS, EG, EO

## Data and materials availability

All data are available in the main text or the supplementary materials.

