## Supplementary material for "Inside-out Signalling From Aminopeptidase N (CD13) To Complement Receptor 3 (CR3, CD11b/CD18)"

**Supplementary Materials**

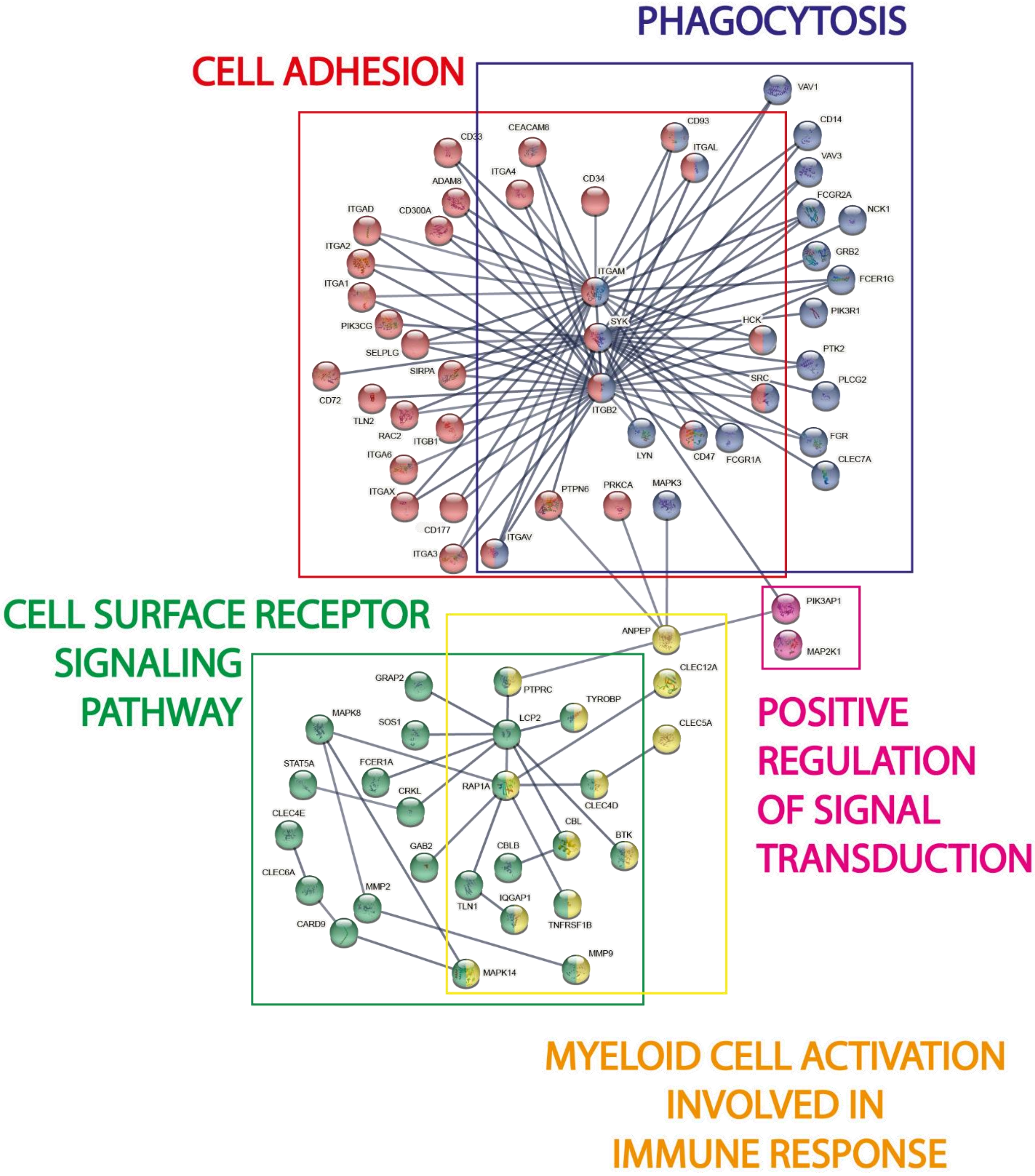

**Fig. S1. Molecular ontology within the protein interaction network.** This diagram depicts the essential functions for which these molecules were included in our interactions network. Each box encloses all the proteins within a category. Cell adhesion and phagocytosis stand out as the main and most specific processes represented by the selected proteins.

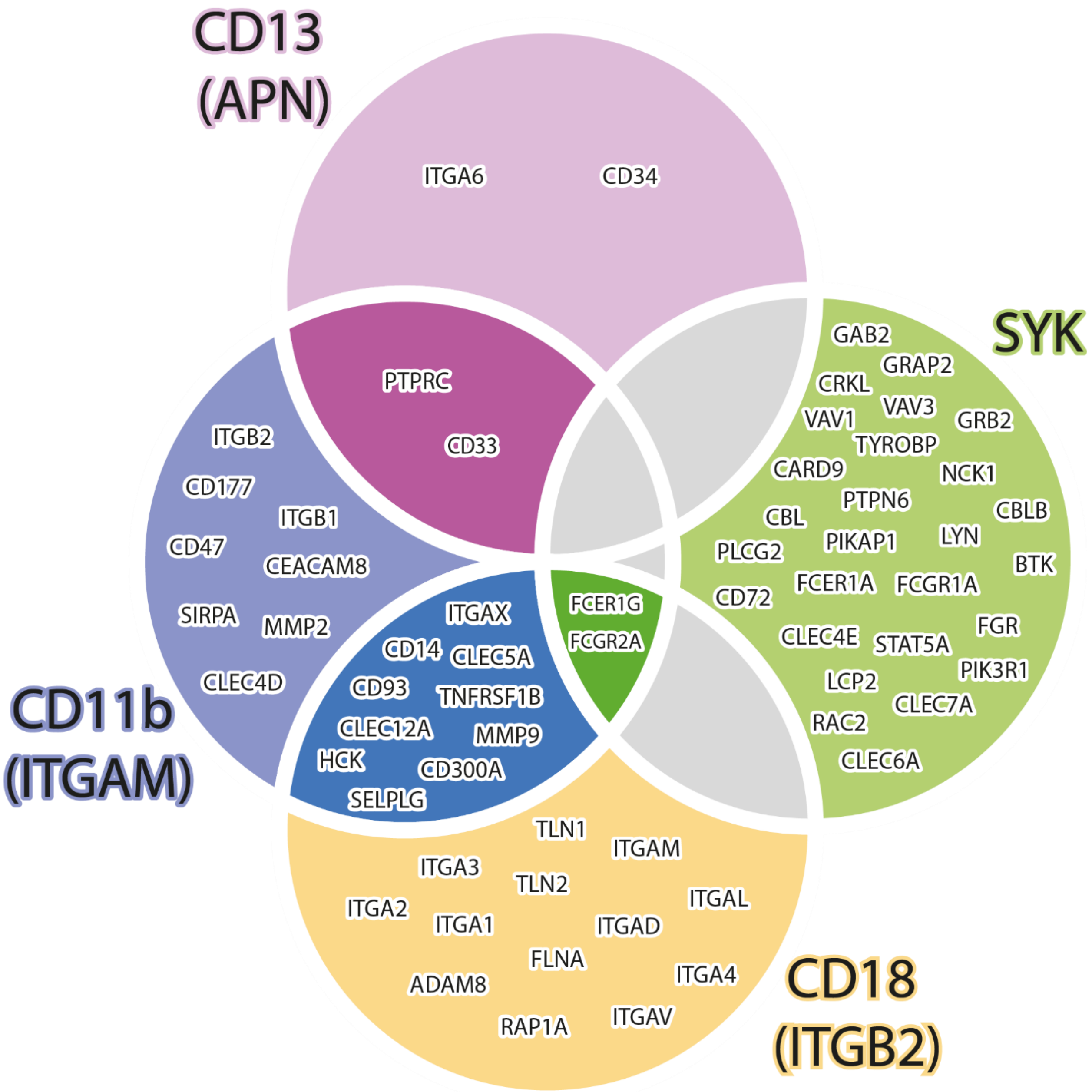

**Fig. S2. 65 non-redundant proteins were deemed of interest after the interrogation of databases using CD13, Syk and CR3 (CD11b/CD18) as queries.** This Venn diagram show each interrogation query as a circle, inside of them are the proteins of interest selected according to our criteria, the molecules that appeared in more than one interrogation/narrowing down process are inside the intersections. The grey areas represent intersections with no common proteins.

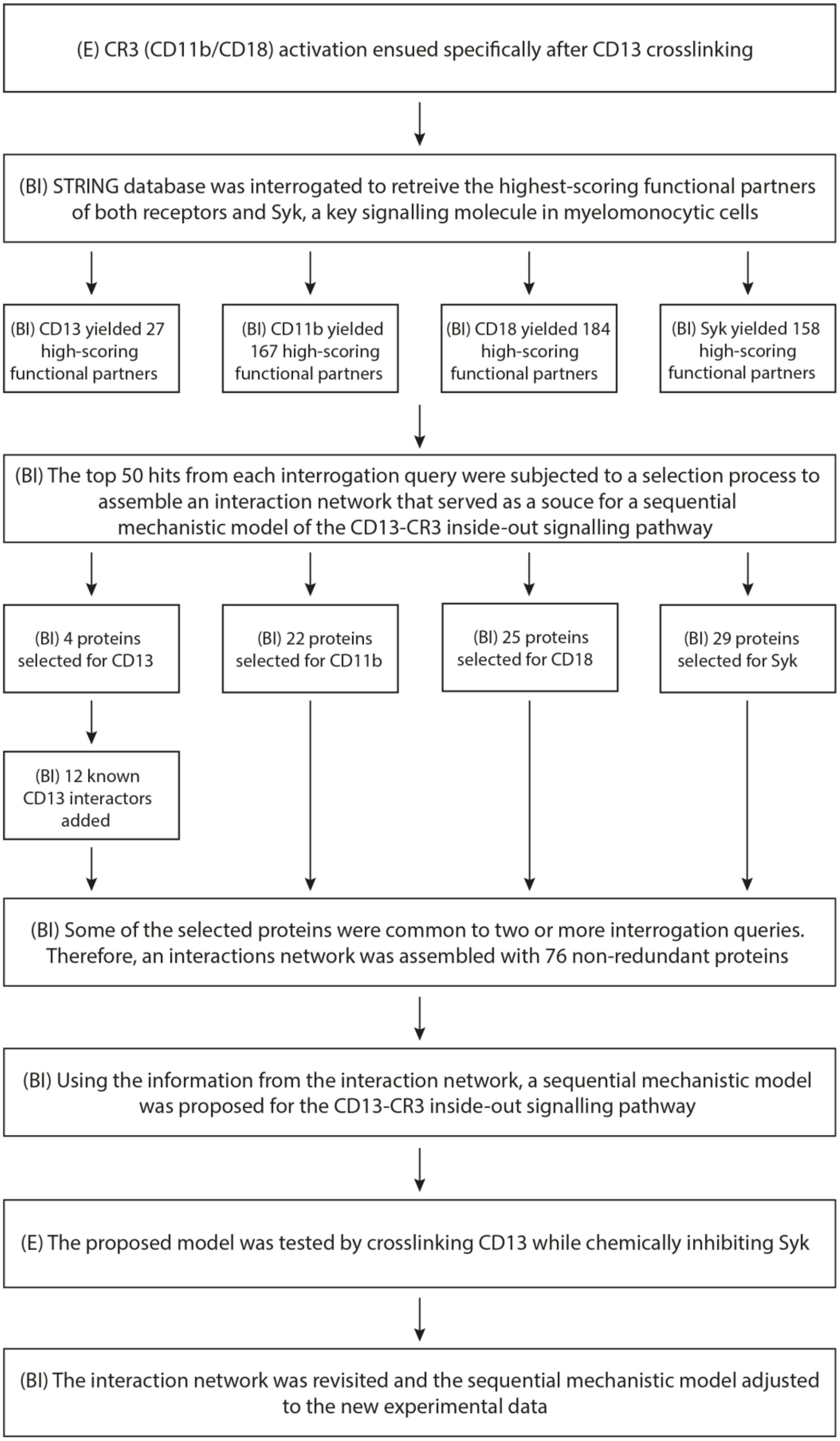
(E) CR3 (CD11b/CD18) activation ensued specifically after CD13 crosslinking
(BI) STRING database was interrogated to retreive the highest-scoring functional partners of both receptors and Syk, a key signalling molecule in myelomonocytic cells
(BI) CD13 yielded 27 high-scoring functional partners
(BI) CD11b yielded 167 high-scoring functional partners
(BI) CD18 yielded 184 high-scoring functional partners
(BI) Syk yielded 158 high-scoring functional partners
(BI) The top 50 hits from each interrogation query were subjected to a selection process to assemble an interaction network that served as a souce for a sequential mechanistic model of the CD13-CR3 inside-out signalling pathway
(BI) 4 proteins selected for CD13
(BI) 22 proteins selected for CD11b
(BI) 25 proteins selected for CD18
(BI) 29 proteins selected for Syk
(BI) 12 known CD13 interactors added
(BI) Some of the selected proteins were common to two or more interrogation queries. Therefore, an interactions network was assembled with 76 non-redundant proteins
(BI) Using the information from the interaction network, a sequential mechanistic model was proposed for the CD13-CR3 inside-out signalling pathway
(E) The proposed model was tested by crosslinking CD13 while chemically inhibiting Syk
(BI) The interaction network was revisited and the sequential mechanistic model adjusted to the new experimental data

**Fig. S3. Workflow.** This flow diagram briefly describes each step taken to tackle the task of proposing a sequential mechanistic model for the newly described CD13-CR3 inside-out signalling pathway. The letter E indicates the experimental steps, whereas the letters BI indicates the bioinformatic ones.

**Table S1. Description of the 76 proteins contained in the CD13, CR3 (CD11b/CD18) and Syk interactions network.** Each node in the protein interactions network (Fig. 3) is enlisted with its Enzime Commission (EC) number, synonyms, function examples (focus on those relevant for our research), corresponding interrogation node and STRING combined score. Data obtained from Gene Cards database, unless otherwise stated. NA = non applicable

| Node | EC number | Synonyms | Functions | Corresponding interrogation node | STRING combined scores |
|---|---|---|---|---|---|
| **ANPEP** | 3.4.11.2 | • CD13<br>• Alanyl Aminopeptidase, Membrane<br>• Gp150 | •Phagocytic receptor(3)<br>•Adhesion Receptor(69)<br>•Myelomonocytic lineage marker(69)<br>•Involved in the processing of various peptides including peptide hormones, neuropeptides, and chemokines<br>•Viral receptor (HCoV-229E, HCMV) | NA – Interrogation node | NA |
| **SYK** | 2.7.10.2 | • p72-Syk<br>• Spleen Associated Tyrosine Kinase | • Non-receptor tyrosine kinase<br>• Inhibits or enhances several biological processes including innate and adaptive immunity, cell adhesion, osteoclast maturation, platelet activation and vascular development | NA – Interrogation node | NA |
| **ITGAM** | NA | • CD11b<br>• CR3 a subunit<br>• Integrin αM | • Phagocytic receptor | Interrogation node / CD18 | NA / 0.992 |

| | | | | | |
|---|---|---|---|---|---|
| | | • Mac-1A<br>• p170 | • Integrin ITGAM/ITGB2 is implicated in various adhesive interactions of monocytes, macrophages, and granulocytes as well as in mediating the uptake of complement-coated particles and pathogens<br>• Mediates neutrophil migration | | |
| **ITGB2** | NA | • CD18<br>• CR3 and CR4 β subunit<br>• Integrin β2<br>• Mac-1 b subunit<br>• p95 | | Interrogation node / CD11b | NA / 0.992 |
| **GRB2** | NA | • Growth Factor Receptor-Bound Protein 2<br>• NCKAP2<br>• ASH | • Adapter protein that provides a critical link between cell surface growth factor receptors and the Ras signaling pathway<br>• Major CBL associated protein | CD13 / Syk | NA / 0.982 |
| **SOS1** | NA | • SOS Ras/Rac Guanine Nucleotide Exchange Factor 1<br>• HGF<br>• GF1<br>• Gingival Fibromatosis, Hereditary, 1<br>• Son Of Sevenless Homolog 1 | • Promotes the exchange of Ras-bound GDP by GTP<br>• Probably by promoting Ras activation, regulates phosphorylation of MAP kinase MAPK3 in response to EGF | CD13 | NA |
| | | • IQ Motif Containing GTPase Activating Protein 1 | • Plays a crucial role in regulating the dynamics and assembly of the actin cytoskeleton. Binds to activated CDC42 but does not stimulate its GTPase activity.<br>• Associates with calmodulin. | | |

| | | | | | |
|---|---|---|---|---|---|
| **IQGAP1** | NA | • p195<br>• HUMORFA01<br>• SAR1<br>• KIAA0051 | • Could serve as an assembly scaffold for the organization of a multimolecular complex that would interface incoming signals to the reorganization of the actin cytoskeleton at the plasma membrane. May promote neurite outgrowth.<br>• May play a possible role in cell cycle by contributing to cell cycle progression after DNA replication arrest | CD13 | NA |
| **PTPRC** | 3.1.3.48 | • Protein Tyrosine Phosphatase Receptor Type C<br>• T200<br>• Gp180<br>• CD45<br>• LCA | • Promiscuous cell surface receptor, it dephosphorylates tyrosines on ITAMs, ITIMs or ITSM son other receptors, regulating their activity. For example, it is capable of inhibiting the anti-phagocytic signal from SIRPA in macrophages, boosting antibody-dependent phagocytosis(*70*) | CD11b / CD13 | 0.97 / 0.814 |
| **CD33** | NA | • SIGLEC-3<br>• Gp67<br>• p67 | • Sialic-acid-binding immunoglobulin-like lectin (Siglec) that plays a role in mediating cell-cell interactions and in maintaining immune cells in a resting state<br>• Src family-mediated phosphorylations on its ITIMs provide docking sites for the recruitment and activation of protein-tyrosine phosphatases PTPN6/SHP-1 and PTPN11/SHP-2<br>• In turn, these phosphatases regulate downstream pathways through | CD13 / CD11b | 0.922 / 0.981 |

| | | | | | |
|---|---|---|---|---|---|
| | | | dephosphorylation of signaling molecules, e.g. phosphoinositide 3-kinase/PI3K | | |
| **CD34** | NA | NA | • Could act as a scaffold for the attachment of lineage specific glycans, allowing stem cells to bind to lectins expressed by stromal cells or other marrow components.<br>• Presents carbohydrate ligands to selectins. | CD13 | 0.867 |
| **ITGA6** | NA | • CD49f<br>• Integrin α6<br>• Integrin α6b<br>• VLA-6 | • ITGA6:ITGB4 binds to NRG1 (via EGF domain) and this binding is essential for NRG1-ERBB signaling<br>• ITGA6:ITGB4 binds to IGF1 and this binding is essential for IGF1 signaling<br>• ITGA6:ITGB4 binds to IGF2 and this binding is essential for IGF2 signaling | CD13 | 0.944 |
| **JNK**(*71*) | 2.7.11.24 | • JNK1: MAPK8, SAPK, JNK-46, PRKM8<br>• JNK2: MAPK9, SAPK1a, JNK-55, PRKM9<br>JNK3: MAPK10, SAPK1b, PRKM10 | • JNKs (c-Jun N-terminal kinases) are a group of mitogen activated protein kinases (MAPKs)<br>• Serine/threonine non-receptor kinases<br>• Signalling pathway effector enzymes, induces by different types of receptors: hormone, neurotransmitters, morphogenic factors, inflammatory cytokines, and those for intracellular and extracellular pathogens<br>• Participate in signalling pathways activated by intracellular stimuli | CD13 | NA |

| | | | | | |
|---|---|---|---|---|---|
| | | | like oxidative stress and DNA damage | | |
| **p38** | 2.7.11.24 | • MAPK11: SAPK2, p38-2, p38b, PRKM11<br><br>• MAPK12: ERK-6, SAPK3, p38g, PRKM12<br><br>• MAPK13: SAPK4, p38d, PRKM13,<br><br>MAPK14: p38a, PRKM14/15, SAPK2a | • Serine/threonine kinase which acts as an essential component of the MAP kinase signal transduction pathway. MAPK14 is one of the four p38 MAPKs which play an important role in the cascades of cellular responses evoked by extracellular stimuli such as proinflammatory cytokines or physical stress leading to direct activation of transcription factors<br>• Phosphorylates the membrane-associated metalloprotease ADAM17 | CD13 | NA |
| **SRC** | 2.7.10.2 | • p60-SRC<br>• SRC1<br>• ASV | • Non-receptor protein tyrosine kinase which is activated following engagement of many different classes of cellular receptors including immune response receptors, integrins and other adhesion receptors, receptor protein tyrosine kinases, G protein-coupled receptors as well as cytokine receptors.<br>• Participates in signalling pathways that control a diverse spectrum of biological activities including gene transcription, immune response, cell adhesion, cell cycle progression, apoptosis, migration, and transformation. | CD13 | NA |

| | | | | | |
|---|---|---|---|---|---|
| | | | • Due to functional redundancy between members of the SRC kinase family, identification of the specific role of each SRC kinase is very difficult.<br>• SRC appears to be one of the primary kinases activated following engagement of receptors and plays a role in the activation of other protein tyrosine kinase (PTK) families.<br>• Receptor clustering or dimerization leads to recruitment of SRC to the receptor complexes where it phosphorylates the tyrosine residues within the receptor cytoplasmic domains. | | |
| **FAK** | 2.7.10.2 | • Protein Tyrosine Kinase 2<br>• PTK2<br>• Focal Adhesion Kinase | • Non-receptor protein-tyrosine kinase that plays an essential role in regulating cell migration, adhesion, spreading, reorganization of the actin cytoskeleton, formation and disassembly of focal adhesions and cell protrusions, cell cycle progression, cell proliferation and apoptosis.<br>• Functions in integrin signal transduction, but also in signalling downstream of numerous growth factor receptors, G-protein coupled receptors (GPCR), EPHA2, netrin receptors and LDL receptors<br>• Promotes activation of MAPK1/ERK2, MAPK3/ERK1 and | CD13 | NA |

| | | | | | |
|---|---|---|---|---|---|
| | | | the MAP kinase signalling cascade | | |
| **ERK 1/2** | 2.7.11.24 | • ERK1: MAPK3, PRKM3, P44-MAPK,<br>• ERK2: MAPK1, PRMK1, P42-MAPk | • Serine/threonine kinase which acts as an essential component of the MAP kinase signal transduction pathway. MAPK1/ERK2 and MAPK3/ERK1 are the 2 MAPKs which play an important role in the MAPK/ERK cascade<br>• Depending on the cellular context, the MAPK/ERK cascade mediates diverse biological functions such as cell growth, adhesion, survival, and differentiation through the regulation of transcription, translation, cytoskeletal rearrangements.<br>• The substrates include protein kinases such as SYK, RAF1, RPS6KA1/RSK1, RPS6KA3/RSK2, RPS6KA2/RSK3, RPS6KA6/RSK4, MKNK1/MNK1, MKNK2/MNK2, RPS6KA5/MSK1, RPS6KA4/MSK2, MAPKAPK3 or MAPKAPK5 | CD13 | NA |
| **PKC** | 2.7.11.13 | • PKCA: PKCa, PRKACA<br>• PKCB: PKCB1/2, PKCb, PRKCB1/2 | • Calcium-activated, phospholipid- and diacylglycerol (DAG)-dependent serine/threonine-protein kinase that is involved in positive and negative regulation of cell proliferation, apoptosis, differentiation, migration and adhesion, tumorigenesis, cardiac | CD13 | NA |

| | | | | | |
|---|---|---|---|---|---|
| | | PKCG: PKCg, PKCC, PRKCG | hypertrophy, angiogenesis, platelet function and inflammation, by directly phosphorylating targets such as RAF1, BCL2, CSPG4, TNNT2/CTNT, or activating signalling cascade involving MAPK1/3 (ERK1/2) and RAP1GAP.<br>• Involved in the stabilization of VEGFA mRNA at post-transcriptional level and mediates VEGFA-induced cell proliferation. In the regulation of calcium-induced platelet aggregation, mediates signals from the CD36/GP4 receptor for granule release, and activates the integrin heterodimer ITGA2B-ITGB3 through the RAP1GAP pathway for adhesion. | | |
| **MEK-1** | 2.7.12.2 | • MAPKK1<br>• MAPK/ERK kinase<br>• ERK activator kinase 1<br>• PRKMK1 | • Dual specificity protein kinase which acts as an essential component of the MAP kinase signal transduction pathway.<br>• Binding of extracellular ligands such as growth factors, cytokines and hormones to their cell-surface receptors activates RAS and this initiates RAF1 activation. RAF1 then further activates the dual-specificity protein kinases MAP2K1/MEK1 and MAP2K2/MEK2. Both MAP2K1/MEK1 and MAP2K2/MEK2 function | CD13 | NA |

| | | | | | |
|---|---|---|---|---|---|
| | | | specifically in the MAPK/ERK cascade and catalyze the concomitant phosphorylation of a threonine and a tyrosine residue in a Thr-Glu-Tyr sequence located in the extracellular signal-regulated kinases MAPK3/ERK1 and MAPK1/ERK2, leading to their activation and further transduction of the signal within the MAPK/ERK cascade. | | |
| **PI3K** (*72*) | 2.7.1.137 | • Phosphatidylinositol-4,5-Bisphosphate 3-Kinase | • There are four Class 1 PI3Ks in mammals (α, β, δ, and γ) that are closely related by sequence homology in the catalytic domain and in the preferential synthesis of $PIP_3$ from $PIP_2$ and ATP<br>• Involved in inflammatory and allergic responses<br>• Modulates chemiotaxis to inflammation sites and in response to chemoattractants<br>• Can control leukocyte polarization and migration through the regulation of spatial PIP3 accumulation, and regulating the organization of F-actin formation and the integrin based on the leading edge<br>• PIK3g (PIKCG) and PIK3d (PIKCD) participate in respiratory burst, chemotaxis and extravasation of neutrophils<br>• PIK3g (PIKCG) and PIK3b | CD13 | NA |

| | | | | | |
|---|---|---|---|---|---|
| | | | (PIKCB) promote platelet aggregation and thrombosis<br>• PIK3g (PIKCG) regulates the adhesive function of integrins αIIb/β3 (ITGA2B/ITGB3) in platelets via a P2Y12 dependent mechanism | | |
| **GAB2** | NA | • GRB2 Associated Binding Protein 2<br>• Pp100<br>• KIAA0571 | • Adapter protein which acts downstream of several membrane receptors including cytokine, antigen, hormone, cell matrix and growth factor receptors to regulate multiple signalling pathways.<br>• In allergic response, it plays a role in mast cells activation and degranulation through PI-3-kinase regulation.<br>• Also involved in the regulation of cell proliferation and haematopoiesis. | Syk | 0.970 |
| **GRAP2** | NA | • Grb2 Related Adaptor Protein 2<br>• Adaptor Protein GRID<br>• Grf-40<br>• Mona<br>• GADS | • Molecular adaptor implied in T-cell activation and macrophage differentiation(73) | Syk | 0.977 |
| **CRKL** | NA | • CRK Like Proto-Oncogene, | • Adaptor molecule asociated with proteins such as WASP, paxilin, Stat-5 and Syk in platelets(40)<br>• Participates, along with Stat-5, in M1 macrophage polarization and in inflammatory response (74) | Syk | 0.981 |

| | | | | | |
|---|---|---|---|---|---|
| | | Adaptor Protein | • Involved in signalling from the negative regulatory receptor CD200R in myeloid cells (74)<br>• Participates in the inhibition of the interferon-mediated proliferation in haematopoietic cells (75) | | |
| **NCK1** | NA | • NCK Adaptor Protein 1<br>• SH2/SH3 Adaptor Protein NCK-Alpha | • Adapter protein which associates with tyrosine-phosphorylated growth factor receptors, such as KDR and PDGFRB, or their cellular substrates. Maintains low levels of EIF2S1 phosphorylation by promoting its dephosphorylation by PP1<br>• May play a role in cell adhesion and migration through interaction with ephrin receptors | Syk | 0.967 |
| **PTPN6** | 3.1.3.48 | • SHP-1<br>• Protein Tyrosine Phosphatase Non-Receptor Type 6<br>• Hematopoietic Cell Protein-Tyrosine Phosphatase<br>• HCP<br>• PTP1C | • Modulates signalling by tyrosine phosphorylated cell surface receptors such as KIT and the EGF receptor/EGFR. The SH2 regions may interact with other cellular components to modulate its own phosphatase activity against interacting substrates<br>• Plays a key role in haematopoiesis | Syk | 0.986 |
| **TYROBP** | NA | • Protein Tyrosine Kinase Binding Protein<br>• DAP12 | • Adapter protein which non-covalently associates with activating receptors found on the surface of a variety of immune cells to mediate signalling and cell activation following ligand binding by the receptors | Syk | 0.988 |

| | | | | | |
|---|---|---|---|---|---|
| | | • KARAP<br>• PLOSL1 | • TYROBP is tyrosine-phosphorylated in the ITAM domain following ligand binding by the associated receptors which leads to activation of additional tyrosine kinases and subsequent cell activation<br>• Also has an inhibitory role in some cells<br>• Non-covalently associates with activating receptors of the CD300 family to mediate cell activation | | |
| **CD72** | NA | LYB2 | • Upon stimulation, it negatively regulates KIT-mediated growth, differentiation, and survival responses in human mast cells (*76*)<br>• Mediates cell adhesion and spreading in murine macrophages via its interaction with soluble SEMA4D/CD100 (*77*)<br>• Soluble CD100 increases *Leishmania amazonensis* promastigotes infection and phagocytosis in murine macrophages in a CD72-dependent fashion (*78*) | Syk | 0.966 |
| **CBL** | 2.3.2.27 | • E3 Ubiquitin-Protein Ligase<br>• Casitas B-Lineage Lymphoma Proto- | • Adapter protein that functions as a negative regulator of many signalling pathways that are triggered by activation of cell surface receptors.<br>• Recognizes activated receptor tyrosine kinases, including KIT, FLT1, FGFR1, FGFR2, PDGFRA, | Syk | 0.993 |

| | | | | | |
|---|---|---|---|---|---|
| | | Oncogene<br>• CBL2 | PDGFRB, CSF1R, EPHA8 and KDR and terminates signalling.<br>• Recognizes membrane-bound HCK, SRC and other kinases of the SRC family and mediates their ubiquitination and degradation.<br>• Participates in signal transduction in haematopoietic cells. | | |
| **CBLB** | 2.3.2.27 | • Cbl Proto-Oncogene B<br>• B-Lineage Lymphoma Proto-Oncogene B<br>• E3 Ubiquitin-Protein Ligase CBL-B | • E3 ubiquitin-protein ligase which accepts ubiquitin from specific E2 ubiquitin-conjugating enzymes, and transfers it to substrates, generally promoting their degradation by the proteasome.<br>• Negatively regulates TCR (T-cell receptor), BCR (B-cell receptor) and FCER1 (high affinity immunoglobulin epsilon receptor) signal transduction pathways.<br>• May be involved in EGFR ubiquitination and internalization | Syk | 0.986 |
| **PIK3R1** | NA | • Phosphoinositide-3-Kinase Regulatory Subunit 1<br>• GRB1<br>• PI3K Subunit p85a<br>• p85-a<br>• IMD36 | • Binds to activated (phosphorylated) protein-Tyr kinases, through its SH2 domain, and acts as an adapter, mediating the association of the p110 catalytic unit to the plasma membrane<br>• Plays a role in ITGB2 signaling<br>• Plays an important role in signaling in response to FGFR1, FGFR2, FGFR3, FGFR4, KITLG/SCF, KIT, PDGFRA and PDGFRB. | Syk | 0.957 |
| | | • Phosphoinositide-3-Kinase Adaptor Protein 1 | • Regulates the inflammatory to reparatory macrophage transition | | |

| | | | | | |
|---|---|---|---|---|---|
| **PIK3AP1** | NA | • B-Cell Phosphoinositide 3-Kinase Adapter Protein 1<br>• BCAP<br>• FLJ35563 | (*79*)<br>• Regulates dendritic cell maturation through the dual-regulation of NF-κB and PI3K/AKT signaling during infection (*80*)<br>• Links TLR signaling to PI3K activation, a process preventing excessive inflammatory cytokine production<br>• Required for macrophage protection from ER stress-induced apoptosis (*81*) | Syk | 0.971 |
| **PLCG2** | 3.1.4.11 | • PLCγ2<br>• Phospholipase C Gamma 2<br>• 1-Phosphatidylinositol 4,5-Bisphosphate Phosphodiesterase Gamma-2<br>• PLC-IV<br>• APLAID<br>• FCAS3 | • The production of the second messenger molecules diacylglycerol (DAG) and inositol 1,4,5-trisphosphate (IP3) is mediated by activated phosphatidylinositol-specific phospholipase C enzymes.<br>• It is a crucial enzyme in transmembrane signalling.<br>• Its function is essential for the CR3-mediated formation of antibacterial extracellular vesicles (*82*) | Syk | 0.991 |
| | | • Lck/Yes-Related Novel Protein Tyrosine Kinase | • Non-receptor tyrosine-protein kinase that transmits signals from cell surface receptors and plays an important role in integrin signalling, the regulation of innate and adaptive immune responses, haematopoiesis, responses to growth factors and cytokines, but also responses to DNA damage and genotoxic agents | | |

| | | | | | |
|---|---|---|---|---|---|
| **LYN** | 2.7.10.2 | • p53Lyn<br>• p56Lyn | • Functions primarily as negative regulator, but can also function as activator, depending on the context<br>• Acts downstream of several immune receptors, including the B-cell receptor, CD79A, CD79B, CD5, CD19, CD22, FCER1, FCGR2, FCGR1A, TLR2 and TLR4 | Syk | 0.966 |
| **FGR** | 2.7.10.2 | • Gardner-Rasheed Feline Sarcoma Viral (V-Fgr) Oncogene Homolog<br>• Src2<br>• p55Fgr<br>• p58Fgr | • Non-receptor tyrosine-protein kinase that transmits signals from cell surface receptors devoid of kinase activity and contributes to the regulation of immune responses, including neutrophil, monocyte, macrophage and mast cell functions, cytoskeleton remodelling in response to extracellular stimuli, phagocytosis, cell adhesion and migration.<br>• Acts downstream of ITGB1 and ITGB2, and regulates actin cytoskeleton reorganization, cell spreading and adhesion<br>• Depending on the context, activates or inhibits cellular responses<br>• Functions as negative regulator of ITGB2 signalling, phagocytosis and SYK activity in monocytes<br>• Required for normal ITGB1 and ITGB2 signalling, normal cell spreading and adhesion in neutrophils and macrophages<br>• Promotes phosphorylation of CBL, | Syk | 0.964 |

| | | | | | |
|---|---|---|---|---|---|
| | | | CTTN, PIK3R1, PTK2/FAK1, PTK2B/PYK2 and VAV2 | | |
| **BTK** | 2.7.10.2 | • Bruton Tyrosine Kinase<br>• B-Cell Progenitor Kinase<br>• BPK<br>• Bruton Agammaglobulinemia Tyrosine Kinase<br>• ATK<br>• AGMX1<br>PSCTK1 | • Tirosina cinasa no receptora indispensable para el desarrollo, diferenciación y señalización en linfocitos B<br>• Non-receptor tyrosine kinase indispensable for B lymphocyte development, differentiation and signalling<br>• BTK acts as a platform to bring together a diverse array of signalling proteins and is implicated in cytokine receptor signalling pathways. Plays an important role in the function of immune cells of innate as well as adaptive immunity, as a component of the Toll-like receptors (TLR) pathway.<br>• The TLR pathway acts as a primary surveillance system for the detection of pathogens and are crucial to the activation of host defence. | Syk | 0.964 |
| | | • C-Type Lectin Domain Family 4 Member E<br>• Macrophage-Inducible C-Type Lectin<br>• C-Type (Calcium | • Calcium-dependent lectin that acts as a pattern recognition receptor (PRR) of the innate immune system: recognizes damage-associated molecular patterns (DAMPs) of abnormal self and pathogen-associated molecular patterns (PAMPs) of bacteria and fungi<br>• Binding of mycobacterial trehalose | | |

| | | | | | |
|---|---|---|---|---|---|
| **CLEC4E** | NA | Dependent, Carbohydrate-Recognition Domain) Lectin, Superfamily Member 9<br>• CLECSF9<br>• MINCLE | 6,6'-dimycolate (TDM) to this receptor complex leads to phosphorylation of the immunoreceptor tyrosine-based activation motif (ITAM) of FCER1G, triggering activation of SYK, CARD9 and NF-kappa-B, consequently driving maturation of antigen-presenting cells and shaping antigen-specific priming of T-cells toward effector T-helper 1 and T-helper 17 cell subtypes | Syk | 0.970 |
| **CLEC7A** | NA | • C-Type Lectin Domain Family 7 Member A<br>• Dectin-1<br>• CLECSF12<br>• SCARE2<br>• CD369<br>• BGR<br>• Dendritic Cell-Associated C-Type Lectin-1<br>• CANDF4 | • Lectin that functions as pattern recognizing receptor (PRR) specific for beta-1,3-linked and beta-1,6-linked glucans, which constitute cell wall constituents from pathogenic bacteria and fungi<br>• Necessary for the TLR2-mediated inflammatory response and activation of NF-kappa-B: upon beta-glucan binding, recruits SYK via its ITAM motif and promotes a signalling cascade that activates some CARD domain-BCL10-MALT1 (CBM) signalosomes, leading to the activation of NF-kappa-B and MAP kinase p38 (MAPK11, MAPK12, MAPK13 and/or MAPK14) pathways which stimulate expression of genes encoding pro-inflammatory cytokines and chemokines | Syk | 0.988 |
| | | • C-Type Lectin Domain | • Calcium-dependent lectin that acts as a PRR of the innate immune | | |

| | | | | | |
|---|---|---|---|---|---|
| **CLEC6A** | NA | Family 6 Member A<br>• Dectin-2<br>• C-Type (Calcium Dependent, Carbohydrate-Recognition Domain) Lectin, Superfamily Member 10<br>• Dendritic Cell-Associated C-Type Lectin 2<br>• CLECSF10 | system: specifically recognizes and binds alpha-mannans on *C. albicans* hyphae<br>• Binding of *C. albicans* alpha-mannans to this receptor complex leads to phosphorylation of the ITAM of FCER1G, triggering activation of SYK, CARD9 and NF-kappa-B, consequently driving maturation of antigen-presenting cells and shaping antigen-specific priming of T-cells toward effector T-helper 1 and T-helper 17 cell subtypes<br>• Up-regulated by granulocyte-macrophage colony-stimulating factor (GM-CSF), TGF-beta 1, TNF-alpha and downregulated by IL-4, IL-10 or UVB in CD14+ monocytes | Syk | 0.971 |
| **CARD9** | NA | • Caspase Recruitment Domain Family Member 9<br>• CANDF2 | • Involved in activation of myeloid cells via classical ITAM-associated receptors and TLR: required for TLR-mediated activation of MAPK, while it is not required for TLR-induced activation of NF-kappa-B<br>• Adapter protein that plays a key role in innate immune response against fungi by forming signalling complexes downstream of C-type lectin receptors | Syk | 0.967 |
| | | | • Adapter protein containing an immunoreceptor tyrosine-based activation motif (ITAM) that | | |

| | | | | | |
|---|---|---|---|---|---|
| **FCER1G** | NA | • Fc Fragment of IgE Receptor Ig<br>• Fc Receptor Gamma-Chain<br>• Fce Receptor Ig<br>• FCRG | transduces activation signals from various immunoreceptors<br>• May function cooperatively with other activating receptors. Functionally linked to integrin beta-2/ITGB2-mediated neutrophil activation<br>• Also involved in integrin alpha-2/ITGA2-mediated platelet activation.<br>• Associates with pattern recognition receptors CLEC4D and CLEC4E to form a functional signalling complex in myeloid cells<br>• Binding of mycobacterial trehalose 6,6'-dimycolate (TDM) to this receptor complex leads to phosphorylation of ITAM, triggering activation of SYK, CARD9 and NF-kappa-B, consequently driving maturation of antigen-presenting cells and shaping antigen-specific priming of T-cells toward effector T-helper 1 and T-helper 17 cell subtypes | Syk / CD11b / CD18 | 0.988 / 0.962 / 0.973 |
| **FCGR2A** | NA | • Fc Fragment of IgG Receptor IIa<br>• CD32<br>• IGFR2<br>• CD32<br>• Low Affinity Immunoglobulin Gamma | • Binds to the Fc region of immunoglobulins gamma. Low affinity receptor<br>• By binding to IgG it initiates cellular responses against pathogens and soluble antigens.<br>• Promotes phagocytosis of opsonized antigens | Syk / CD11b / CD18 | 0.974 / 0.987 / 0.947 |

| | | | | | |
|---|---|---|---|---|---|
| | | Fc Region Receptor II-A | | | |
| **FCGR1A** | NA | • Fc Fragment of IgG Receptor Ia<br>• High Affinity Immunoglobulin Gamma Fc Receptor I<br>• CD64A<br>• IGFR1 | • High affinity receptor for the Fc region of immunoglobulins gamma<br>• Functions in both innate and adaptive immune responses | Syk | 0.976 |
| **FCER1A** | NA | • High Affinity Immunoglobulin Epsilon Receptor Subunit Alpha<br>• FCE1A<br>• FcER1 | • Binds to the Fc region of immunoglobulins epsilon<br>• High affinity receptor responsible for initiating the allergic response. Binding of allergen to receptor-bound IgE leads to cell activation and the release of mediators (such as histamine) responsible for the manifestations of allergy<br>• The same receptor also induces the secretion of important lymphokines. | Syk | 0.963 |
| **LCP2** | NA | • Lymphocyte Cytosolic Protein 2<br>• SH2 Domain-Containing Leukocyte Protein Of 76 KDa<br>• SLP76 | • Diseases associated with adaptor protein LCP2 include Immunodeficiency 81 and Wiskott-Aldrich Syndrome<br>• When occupied, $\alpha_v\beta_3$ on osteoclasts activates a canonical signaling complex consisting of c-Src, Syk, Dap12, Slp76, Vav 3, and Rac that permits the cell to spread and form actin rings(83) | Syk | 0.994 |
| | | | • Alongside STAT5B and STAT3, regulates the balance of pro- and | | |

| | | | | | |
|---|---|---|---|---|---|
| **STAT5A** | NA | • Signal Transducer and Activator of Transcription 5A<br>• MGF<br>• Epididymis Secretory Sperm Binding Protein | anti-inflammatory cytokines in PRR-stimulated macrophages (*84*)<br>• Carries out a dual function: signal transduction and activation of transcription<br>• May mediate cellular responses to activated FGFR1, FGFR2, FGFR3 and FGFR4<br>• Regulates the expression of milk proteins during lactation. | Syk | 0.968 |
| **VAV1** | NA | • Vav Guanine Nucleotide Exchange Factor 1 | • Couples tyrosine kinase signals with the activation of the Rho/Rac GTPases, thus leading to cell differentiation and/or proliferation<br>• VAV proto-oncogene 1,homolog,expressed in haematopoietic cells, critical transducer of T cell receptor signals to the calcium, ERK and FNKB pathways | Syk | 0.999 |
| **VAV3** | NA | • Vav Guanine Nucleotide Exchange Factor 3 | • Exchange factor for GTP-binding proteins RhoA, RhoG and, to a lesser extent, Rac1.<br>• Binds physically to the nucleotide-free states of those GTPases.<br>• Responsible for integrin beta-2 (ITGB2)-mediated macrophage adhesion and, to a lesser extent, contributes to beta-3 (ITGB3)-mediated adhesion. Does not affect integrin beta-1 (ITGB1)-mediated adhesion<br>• May be important for integrin-mediated signalling, at least in some | Syk | 0.986 |

| | | | | | |
|---|---|---|---|---|---|
| | | | cell types. In osteoclasts, along with SYK tyrosine kinase, required for signalling through integrin alpha-v/beta-1 (ITAGV-ITGB1), a crucial event for osteoclast proper cytoskeleton organization and function.<br>• Necessary for proper wound healing. In the course of wound healing, required for the phagocytotic cup formation preceding macrophage phagocytosis of apoptotic neutrophils. | | |
| **RAC2** | NA | • Rac Family Small GTPase 2<br>• Ras-Related C3 Botulinum Toxin Substrate 2<br>• p21-rac2<br>• EN-7<br>• HSPC022<br>• GX | • Plasma membrane-associated small GTPase which cycles between an active GTP-bound and inactive GDP-bound state<br>• In active state binds to a variety of effector proteins to regulate cellular responses, such as secretory processes, phagocytose of apoptotic cells and epithelial cell polarization.<br>• Augments the production of reactive oxygen species (ROS) by NADPH oxidase. | Syk | 0.957 |
| **ITGB1** | NA | • Integrin β1<br>• Fibronectin Receptor Subunit Beta<br>• Very Late Activation Protein, Beta Polypeptide<br>• CD29 | • Integrins alpha-1/beta-1, alpha-2/beta-1, alpha-10/beta-1 and alpha-11/beta-1 are receptors for collagen. Integrins alpha-1/beta-1 and alpha-2/beta-2 recognize the proline-hydroxylated sequence G-F-P-G-E-R in collagen<br>• Beta-1 integrins recognize the sequence R-G-D in a wide array of | CD11b | 0.96 |

| | | | | | |
|---|---|---|---|---|---|
| | | • MDF2<br>• MSK12<br>• Glycoprotein IIa<br>• FNRB | ligands. When associated with alpha-7 integrin, regulates cell adhesion and laminin matrix deposition. Involved in promoting endothelial cell motility and angiogenesis | | |
| **ITGAX** | NA | • CD11c<br>• Integrin Subunit Alpha X<br>• Integrin, Alpha X (Complement Component 3 Receptor 4 Subunit)<br>• Leu M5<br>• SLEB6<br>• Leukocyte Surface Antigen P150,95, Alpha Subunit | • Integrin alpha X cell surface adhesion receptor mediating cell-adhesion to extra cellular matrix or to other cells, through hetero dimerization and connecting to the cytoskeleton and various signalling molecules within cells, dimerizing with ITGB2 in fibrinogen, C3b receptor | CD11b / CD18 | 0.954 / 0.997 |
| **CEACAM8** | NA | • CEA Cell Adhesion Molecule 8<br>• Carcinoembryonic Antigen-Related Cell Adhesion Molecule 8<br>• CD67<br>• CD66b<br>• CGM6<br>• NCA-95 | • Cell surface glycoprotein that plays a role in cell adhesion in a calcium-independent manner<br>• Mediates heterophilic cell adhesion with other carcinoembryonic antigen-related cell adhesion molecules, such as CEACAM6<br>• Heterophilic interaction with CEACAM8 occurs in activated neutrophils | CD11b | 0.969 |
| | | | • Coreceptor for bacterial lipopolysaccharide<br>• Acts via MyD88, TIRAP and TRAF6, leading to NF-kappa-B activation, cytokine secretion and the inflammatory response | | |

| | | | | | |
|---|---|---|---|---|---|
| **CD14** | NA | Myeloid Cell-Specific Leucine-Rich Glycoprotein | • Acts as a coreceptor for TLR2:TLR6 heterodimer in response to diacylated lipopeptides and for TLR2:TLR1 heterodimer in response to triacylated lipopeptides, these clusters trigger signalling from the cell surface and subsequently are targeted to the Golgi in a lipid-raft dependent pathway | CD11b / CD18 | 0.985 / 0.979 |
| **CD93** | NA | • C1qR(P)<br>• CDw93<br>• Complement Component 1 Q Subcomponent Receptor 1<br>• Matrix-Remodeling-Associated Protein 4<br>• MXRA4<br>• ECSM3<br>• DJ737E23.1 | • Receptor (or element of a larger receptor complex) for C1q, mannose-binding lectin (MBL2) and pulmonary surfactant protein A (SPA)<br>• May mediate the enhancement of phagocytosis in monocytes and macrophages upon interaction with soluble defence collagens<br>• May play a role in intercellular adhesion | CD11b / CD18 | 0.962 / 0.936 |
| **CD300A** | NA | • Immunoglobulin Superfamily Member 12<br>• IGSF12<br>• CMRF35-Like Molecule 8<br>• CMRF-35-H9<br>• CMRF5H<br>• NK Inhibitory Receptor<br>• IRC1/IRC2 | • Inhibitory receptor which may contribute to the downregulation of cytolytic activity in natural killer (NK) cells, and to the downregulation of mast cell degranulation<br>• Negatively regulates the Toll-like receptor (TLR) signalling mediated by MYD88 but not TRIF through activation of PTPN6 | CD11b / CD18 | 0.951 / 0.949 |
| | | • HNA2a | • In association with beta-2 integrin heterodimer ITGAM/CD11b and | | |

| | | | | | |
|---|---|---|---|---|---|
| **CD177** | | • PRV1<br>• NB1<br>• Polycythemia Rubra Vera Protein 1<br>• Human Neutrophil Alloantigen 2a | ITGB2/CD18, mediates activation of TNF-alpha primed neutrophils including degranulation and superoxide production<br>• In addition, by preventing beta-2 integrin internalization and attenuating chemokine signaling favors adhesion over migration<br>• By displaying PRTN3 at the neutrophil cell surface, may play a role in enhancing endothelial cell junctional integrity and thus vascular integrity during neutrophil diapedesis | CD11b | 0.937 |
| **CD47** | NA | • Integrin-Associated Signal Transducer<br>• IAP<br>• MER6<br>• OA3 | • Has a role in both cell adhesion by acting as an adhesion receptor for THBS1 on platelets, and in the modulation of integrins<br>• Receptor for SIRPA, binding to which prevents maturation of immature dendritic cells and inhibits cytokine production by mature dendritic cells<br>• Interaction with SIRPG mediates cell-cell adhesion, enhances superantigen-dependent T-cell-mediated proliferation and costimulates T-cell activation<br>• May play a role in membrane transport and/or integrin dependent signal transduction | CD11b | 0.95 |
| | | | • Immunoglobulin-like cell surface receptor for CD47. Acts as docking protein and induces translocation of | | |

| | | | | | |
|---|---|---|---|---|---|
| **SIRPA** | NA | • Signal Regulatory Protein Alpha<br>• SHPS1<br>• BIT<br>• MFR<br>• p84<br>• CD172a<br>• MYD-1 | PTPN6, PTPN11 and other binding partners from the cytosol to the plasma membrane<br>• Supports adhesion of cerebellar neurons, neurite outgrowth and glial cell attachment<br>• Involved in the negative regulation of receptor tyrosine kinase-coupled cellular responses induced by cell adhesion, growth factors or insulin<br>• Mediates negative regulation of phagocytosis, mast cell activation and dendritic cell activation<br>• CD47 binding prevents maturation of immature dendritic cells and inhibits cytokine production by mature dendritic cells | CD11b | 0.96 |
| **HCK** | 2.7.10.2 | • Hemopoietic Cell Kinase<br>• p59-HCK/p60-HCK<br>• JTK9 | • Non-receptor tyrosine-protein kinase found in hematopoietic cells that transmits signals from cell surface receptors and plays an important role in the regulation of innate immune responses, including neutrophil, monocyte, macrophage and mast cell functions, phagocytosis, cell survival and proliferation, cell adhesion and migration<br>• Acts downstream of integrins, such as ITGB1 and ITGB2, and receptors that bind the Fc region of immunoglobulins, such as FCGR1A and FCGR2A, but also CSF3R, PLAUR, the receptors for IFNG, | CD11b / CD18 | 0.965 / 0.979 |

| | | | | | |
|---|---|---|---|---|---|
| | | | IL2, IL6 and IL8.<br>• During the phagocytic process, mediates mobilization of secretory lysosomes, degranulation, and activation of NADPH oxidase to bring about the respiratory burst<br>• Plays a role in the release of inflammatory molecules<br>• Promotes reorganization of the actin cytoskeleton and actin polymerization, formation of podosomes and cell protrusions<br>• Phosphorylates CBL in response to activation of immunoglobulin gamma Fc region receptors. Phosphorylates ADAM15, BCR, ELMO1, FCGR2A, GAB1, GAB2, RAPGEF1, STAT5B, TP73, VAV1 and WAS | | |
| **MMP2** | 3.4.24.24 | • Gelatinase A<br>• Matrix Metallopeptidase 2<br>• TBE-1<br>• 72 KDa Type IV Collagenase<br>• CLG4A<br>• MONA | • Ubiquitinous metalloproteinase that is involved in diverse functions such as remodeling of the vasculature, angiogenesis, tissue repair, tumor invasion, inflammation, and atherosclerotic plaque rupture<br>• As well as degrading extracellular matrix proteins, can also act on several nonmatrix proteins such as big endothelial 1 and beta-type CGRP promoting vasoconstriction<br>• PEX, the C-terminal non-catalytic fragment of MMP2, posseses anti-angiogenic and anti-tumor properties and inhibits cell | CD11b | 0.941 |

| | | | | | |
|---|---|---|---|---|---|
| | | | migration and cell adhesion to FGF2 and vitronectin. Ligand for integrinv/beta3 on the surface of blood vessels | | |
| **MMP9** | 3.4.24.35 | • Gelatinase B<br>• GELB<br>• Matrix Metallopeptidase 9<br>• 92 KDa Type IV Collagenase<br>• CLG4B | • Matrix metalloproteinase that plays an essential role in local proteolysis of the extracellular matrix and in leukocyte migration<br>• Up-regulated by ARHGEF4, SPATA13 and APC via the JNK signalling pathway in colorectal tumour cells | CD11b / CD18 | 0.969 / 0.946 |
| **CLEC4D** | NA | • C-Type Lectin Domain Family 4 Member D<br>• C-Type (Calcium Dependent, Carbohydrate-Recognition Domain) Lectin, Superfamily Member 8<br>• C-Type Lectin-Like Receptor 6<br>• MCL<br>• Dectin-3<br>CD368 | • Calcium-dependent lectin that acts as a pattern recognition receptor (PRR) of the innate immune system: recognizes damage-associated molecular patterns (DAMPs) of pathogen-associated molecular patterns (PAMPs) of bacteria and fungi<br>• Interacts with signalling adapter Fc receptor gamma chain/FCER1G, likely via CLEC4E, to form a functional complex in myeloid cells<br>• Binding of mycobacterial TDM or *C. albicans* alpha-mannans to this receptor complex leads to phosphorylation of the immunoreceptor tyrosine-based activation motif (ITAM) of FCER1G, triggering activation of SYK, CARD9 and NF-kappa-B, consequently driving maturation of antigen-presenting cells and shaping | CD11b | 0.943 |

| | | | | | |
|---|---|---|---|---|---|
| | | | antigen-specific priming of T-cells toward effector T-helper 1 and T-helper 17 cell subtypes | | |
| **CLEC12A** | NA | • C-Type Lectin Domain Family 12 Member A<br>• Myeloid Inhibitory C-Type Lectin-Like Receptor<br>• Dendritic Cell-Associated Lectin 2<br>• DCAL-2<br>• CLL-1<br>• MICL<br>• CD371 | • Cell surface receptor that modulates signalling cascades and mediates tyrosine phosphorylation of target MAP kinases<br>• Downregulated in activated leukocytes recruited to a site of inflammation | CD11b / CD18 | 0.950 / 0.933 |
| **CLEC5A** | NA | • C-Type Lectin Domain Containing 5[a]<br>• MDL-1<br>• Myeloid DAP12-Associating Lectin-1 | • Cell surface receptor that signals via TYROBP<br>• Regulates inflammatory responses<br>• Critical macrophage receptor for dengue virus serotypes 1-4<br>• The binding of dengue virus to CLEC5A triggers signalling through the phosphorylation of TYROBP. This interaction does not result in viral entry, but stimulates proinflammatory cytokine release | CD11b / CD18 | 0.947 / 0.934 |
| **SELPLG** | NA | • CLA<br>• Selectin P Ligand<br>• PSLG-1<br>• CD162 | • A SLe(x)-type proteoglycan, which through high affinity, calcium-dependent interactions with E-, P- and L-selectins, mediates rapid rolling of leukocytes over vascular surfaces during the initial steps in inflammation<br>• Critical for the initial leukocyte | CD11b / CD18 | 0.962 / 0.966 |

| | | | | | |
|---|---|---|---|---|---|
| | | | capture<br>• Acts as a receptor for enterovirus 71 | | |
| **TNFRSF1B** | NA | • CLA<br>• TNF Receptor Superfamily Member 1B<br>• TNFBR<br>• P75<br>• TNF-R75<br>• TNF-RII<br>• CD120b<br>• TNFR80<br>• TNFR2 | • Receptor with high affinity for TNFSF2/TNF-alpha and approximately 5-fold lower affinity for homotrimeric TNFSF1/lymphotoxin-alpha. The TRAF1/TRAF2 complex recruits the apoptotic suppressors BIRC2 and BIRC3 to TNFRSF1B/TNFR2<br>• This receptor mediates most of the metabolic effects of TNF-alpha<br>• Isoform 2 blocks TNF-alpha-induced apoptosis, which suggests that it regulates TNF-alpha function by antagonizing its biological activity | CD11b / CD18 | 0.948 / 0.939 |
| **ITGAL** | NA | • CD11a<br>• Integrin Subunit Alpha L<br>• Lymphocyte Function-Associated Antigen 1, Alpha Polypeptide<br>• LFA-1A<br>• P180 | • Integrin ITGAL/ITGB2 is a receptor for ICAM1, ICAM2, ICAM3 and ICAM4<br>• Involved in a variety of immune phenomena including leukocyte-endothelial cell interaction, cytotoxic T-cell mediated killing, and antibody dependent killing by granulocytes and monocytes<br>• Contributes to natural killer cell cytotoxicity<br>• Involved in leukocyte adhesion and transmigration of leukocytes including T-cells and neutrophils<br>• Integrin ITGAL/ITGB2 in association with ICAM3, contributes to apoptotic neutrophil | CD18 | 0.999 |

| | | | | | |
|---|---|---|---|---|---|
| | | | phagocytosis by macrophages | | |
| **ITGAD** | NA | • Integrin Subunit Alpha D<br>• CD11d<br>• ADB2 | • Cell surface adhesion receptor mediating cell-adhesion to extra cellular matrix or to other cells, through hetero dimerization and connecting to the cytoskeleton and various signalling molecules within cells, arrayed in tandem with ITGAX (CD11C),dimerizing with ITGB2 in fibrinogen,C3b receptor<br>• Receptor for ICAM3 and VCAM1<br>• May play a role in the atherosclerotic process such as clearing lipoproteins from plaques and in phagocytosis of blood-borne pathogens, particulate matter, and senescent erythrocytes from the blood | CD18 | 0.983 |
| **ITGA4** | NA | • Integrin Subunit Alpha 4<br>• CD49d | • Integrin, alpha 4, cell surface adhesion receptor mediating cell-adhesion to extra cellular matrix or to other cells, through hetero dimerization and connecting to the cytoskeleton and various signalling molecules within cells, component of VLA-4 receptor, dimerizing with ITGB1 or ITGB7 in fibronectin, VCAM1 receptors<br>• Integrin alpha-4/beta-7 is also a receptor for MADCAM1<br>• It recognizes the sequence L-D-T in MADCAM1. On activated endothelial cells integrin VLA-4 triggers homotypic aggregation for | CD18 | 0.983 |

| | | | | | |
|---|---|---|---|---|---|
| | | | most VLA-4-positive leukocyte cell lines | | |
| **ITGAV** | NA | • Integrin Subunit Alpha V<br>• Vitronectin Receptor Subunit Alpha<br>• CD51<br>• MSK8<br>• VNRA<br>• VTNR | • The alpha-V (ITGAV) integrins are receptors for vitronectin, cytotactin, fibronectin, fibrinogen, laminin, matrix metalloproteinase-2, osteopontin, osteomodulin, prothrombin, thrombospondin and vWF<br>• ITGAV:ITGB5 acts as a receptor for Adenovirus type C<br>• ITGAV:ITGB3 acts as a receptor for Herpes virus 8/HHV-8 | CD18 | 0.971 |
| **ITGA2** | NA | • Integrin Subunit Alpha 2<br>• CD49b<br>• Very Late Activation Protein 2 Receptor, Alpha-2 Subunit<br>• GPIa<br>• HPA-5 | • Integrin alpha-2/beta-1 is a receptor for laminin, collagen, collagen C-propeptides, fibronectin and E-cadherin<br>• It is responsible for adhesion of platelets and other cells to collagens, modulation of collagen and collagenase gene expression, force generation and organization of newly synthesized extracellular matrix<br>• Integrin ITGA2:ITGB1 acts as a receptor for Human rotavirus A | CD18 | 0.956 |
| **ITGA3** | NA | • Integrin Subunit Alpha 3<br>• CD49c<br>• Alpha 3 Subunit Of VLA-3 Receptor<br>• Galactoprotein B3<br>• GAP-B3 | • Integrin alpha-3/beta-1 is a receptor for fibronectin, laminin, collagen, epiligrin, thrombospondin and CSPG4<br>• Integrin alpha-3/beta-1 provides a docking site for FAP (seprase) at invadopodia plasma membranes in a collagen-dependent manner and | CD18 | 0.969 |

| | | | | | |
|---|---|---|---|---|---|
| | | • FRP-2<br>• MSK18 | hence may participate in the adhesion, formation of invadopodia and matrix degradation processes, promoting cell invasion<br>• Alpha-3/beta-1 may mediate with LGALS3 the stimulation by CSPG4 of endothelial cells migration | | |
| **ITGA1** | NA | • Integrin Subunit Alpha 1<br>• CD49a<br>• Alpha 1 Subunit Of VLA-3 Receptor<br>• Laminin And Collagen Receptor | • Cell surface adhesion receptor mediating cell-adhesion to extra cellular matrix or to other cells, through hetero dimerization and connecting to the cytoskeleton and various signalling molecules within cells, dimerizing with ITGB1 in collagen, laminin receptors | CD18 | 0.960 |
| **ADAM8** | 3.4.24.- | • Disintegrin And Metalloproteinase Domain-Containing Protein 8<br>• Cell Surface Antigen MS2<br>• CD156 | • A disintegrin and metalloprotease (active) domain 8, membrane anchored cell surface adhesion protein (antigen MS2), expressed in granulocyte, monocyte, macrophage, involved in cell-cell and cell-matrix interactions<br>• Possible involvement in extravasation of leukocytes | CD18 | 0.954 |
| **RAP1A** | 3.6.5.2 | • Rap1<br>• RAP1A, Member Of RAS Oncogene Family<br>• KREV-1<br>• SMGP21<br>• C21KG | • Induces morphological reversion of a cell line transformed by a Ras oncogene. Counteracts the mitogenic function of Ras, at least partly because it can interact with Ras GAPs and RAF in a competitive manner. Together with ITGB1BP1, regulates KRIT1 localization to microtubules and membranes | CD18 | 0.936 |

| | | | | | |
|---|---|---|---|---|---|
| | | • G-22K | | | |
| **TLN1** | NA | • Talin-1<br>• ILWEQ<br>• KIAA1027 | • Talin-1 contributes to alpha(4)beta(1)-dependent chemotaxis, suggesting that it participates in a later stage of the leukocyte adhesion cascade when the leukocyte cytoskeleton undergoes dramatic rearrangement (*85*)<br>• Overexpression of the talin-1 head domain results in separation of LFA-1 cytoplasmic tails, which is also observed following chemokine stimulation and correlates with integrin conformational change and activation (*86*) | CD18 | 0.98 |
| **TLN2** | NA | • Talin-2<br>• ILWEQ<br>• KIAA0320 | • Human macrophages express both Talin isoforms, 1 and 2(*87*)<br>• *Tln2* seems to be the ancestral gene, Tln1 would appear by gene duplication early in the cordate lineage (*88*)<br>• As a major component of focal adhesion plaques that links integrin to the actin cytoskeleton, may play an important role in cell adhesion | CD18 | 0.951 |
| **FLNA** | 2.1.1.43<br>6.3.4.4 | • Filamin A<br>• ABP-280<br>• Endothelial Actin-Binding Protein | • Promotes orthogonal branching of actin filaments and links actin filaments to membrane glycoproteins. Anchors various transmembrane proteins to the actin cytoskeleton and serves as a scaffold | CD18 | 0.965 |

| | | | |
|---|---|---|---|
| | | • ABPX<br>• CSBS<br>• CVD1<br>• FGS2<br>• NHBP<br>• OPD1/2 | for a wide range of cytoplasmic signalling proteins<br>• Plays a role in cell-cell contacts and adherent junctions during the development of blood vessels, heart and brain organs. Plays a role in platelets morphology through interaction with SYK that regulates ITAM- and ITAM-like-containing receptor signalling, resulting in by platelet cytoskeleton organization maintenance |
